# Cross-validation of distance measurements in proteins by PELDOR/DEER and single-molecule FRET

**DOI:** 10.1101/2020.11.23.394080

**Authors:** Martin F. Peter, Christian Gebhardt, Rebecca Mächtel, Janin Glaenzer, Gavin H. Thomas, Thorben Cordes, Gregor Hagelueken

## Abstract

Pulsed electron-electron double resonance spectroscopy (PELDOR or DEER) and single molecule Förster resonance energy transfer spectroscopy (smFRET) are recent additions to the toolbox of integrative structural biology. Both methods are frequently used to visualize conformational changes and to determine nanometer-scale distances in biomacromolecules including proteins and nucleic acids. A prerequisite for the application of PELDOR/DEER and smFRET is the presence of suitable spin centers or fluorophores in the target molecule, which are usually introduced via chemical biology methods. The application portfolio of the two methods is overlapping: each allows determination of distances, to monitor distance changes and to visualize conformational heterogeneity and -dynamics. Both methods can provide qualitative information that facilitates mechanistic understanding, for instance on conformational changes, as well as quantitative data for structural modelling. Despite their broad application, a comprehensive comparison of the accuracy of PELDOR/DEER and smFRET is still missing and we set out here to fill this gap. For this purpose, we prepared a library of double cysteine mutants of three well-studied substrate binding proteins that undergo large-scale conformational changes upon ligand binding. The distances between the introduced spin- or fluorescence labels were determined via PELDOR/DEER and smFRET, using established standard experimental protocols and data analysis routines. The experiments were conducted in the presence and absence of the natural ligands to investigate how well the ligand-induced conformational changes could be detected by the two methods. Overall, we found good agreement for the determined distances, yet some surprising inconsistencies occurred. In our set of experiments, we identified the source of discrepancies as the use of cryoprotectants for PELDOR/DEER and label-protein interactions for smFRET. Our study highlights strength and weaknesses of both methods and paves the way for a higher confidence in quantitative comparison of PELDOR/DEER and smFRET results in the future.

## Introduction

Since the determination of the first macromolecular structures in the 1950s, our knowledge of the structure and function of macromolecules has dramatically increased. At the time of writing, the protein database (PDB, www.rcsb.org) contained more than 170,000 structures. Many of these PDB entries represent identical macromolecules in different conformations. This might reflect varying experimental conditions or illustrate the dynamic nature of proteins, i.e., the large- or small-scale structural fluctuations that are crucial aspects of their biological function. ABC-transporters are just one prominent example ^1,2^: Without the ability of these membrane proteins to perform large and controlled conformational changes, efficient active solute transport across membranes would be impossible. On a smaller distance scale also enzymes require structural dynamics, be it for their catalytic activity per se or for its regulation ^3,4^.

Until now, most available macromolecular structures were determined by X-ray crystallography, ~10 % by NMR, and ~2 % by cryo-EM (https://www.rcsb.org). Undoubtedly, cryo-EM and X-ray crystallography can deliver highly detailed insights into the molecular scaffolds of proteins. Nevertheless, they have the disadvantage that such structures are not determined in liquid solution, but in a crystal lattice or frozen on an EM grid. The underlying macromolecular dynamics can often only be inferred by determining multiple structures and combining them into a molecular “movie”^5^. Such movies require additional (biochemical) support to verify the selected order of structural states. Traditionally, the study of such dynamic processes is the strength of nuclear magnetic resonance spectroscopy (NMR). But, this method is limited to the study of relatively small proteins (typically <70 kDa; larger homo-oligomers are an exception), which renders the analysis of many proteins impossible. Due to the well-known limitations of the three major players, “integrative” methods have become increasingly popular in the last decade. The idea behind this concept is to combine models from either of the three mentioned approaches with data from e.g., hydrogen-exchange mass spectrometry HDX-MS ^6^, Förster resonance energy transfer FRET ^7–10^, small angle X-ray scattering SAXS ^11^ or pulsed electron-electron double resonance spectroscopy PELDOR (also known as double electron-electron resonance spectroscopy, DEER) ^12^. These orthogonal techniques allow to study conformational dynamics, to visualize conformational heterogeneity, to derive distance constrains between selected residues, and to determine entire contact interfaces even for heterogenous samples in a near-physiological environment. Such information is very hard or even impossible to obtain with the classical structural biology techniques alone.

In this study, we compared two popular methods to determine distances in proteins: single-molecule FRET (smFRET) and PELDOR/DEER spectroscopy. Both techniques are suitable to determine interprobe distances at the nanometer scale and thereby to detect conformational changes of macromolecules in the (frozen) solution state. We conducted a cross-validation of the two techniques using three model protein systems: (i) HiSiaP, the periplasmic substrate binding protein (SBP) from the sialic acid TRAP transporter of *Haemophilus influenzae*^13,14^, (ii) MalE, also known as MBP (maltose binding protein) from *Escherichia coli*, which plays an important role in the uptake of maltose and maltodextrins by the maltose transporter, MalEFGK2 ^15,16^ and (iii) SBD2, the second of two substrate binding domains that are constituents of the glutamine ABC transporter GlnPQ from *Lactococcus lactis*^17–20^. According to published studies, these SBPs have a common ligand binding mechanism but fall within distinct structural sub categories of substrate-binding proteins ^21^. It is also well established that upon binding of substrate, SBPs undergo a large conformational shift (> 10 Å for selected residues) from an open unliganded conformation (apo) to a closed conformation (holo) (Figure 1, A) ^21–23^. The conformational landscape of SBPs has now been studied for more than four decades using a variety of techniques including crystallography, energy calculations, simulations, NMR, PELDOR/DEER and smFRET ^6,16,17,20,24–33^. Importantly, crystal structures for substrate-free and substrate-bound conformation, exist for all our target proteins. This enabled us to build structural models of the labelled protein complexes and helped us to interpret the measured distances. Moreover, substrate binding to these proteins has been investigated and quantified in previous studies ^13,17,24,31,34–36^. Finally, the selected SBPs are small, soluble and relatively easy to produce in *E. coli*. The proteins are therefore well suited for our purpose, i.e., to apply PELDOR/DEER and smFRET standard procedures for distance determination and then to objectively compare the results. The two methods have only rarely been directly pitted against each other on the same macromolecular systems using identical labelling sites ^37–39^ and to date, a systematic comparison of the two methods is lacking. Since PELDOR/DEER and smFRET are often independently used to validate structural models, a cross validation is important to objectively judge their accuracy, to gauge the severity of their distinct limitations, to detect possible further pitfalls or just to choose the most suitable method for distance experiments in a biological system of interest.

**Figure 1:**
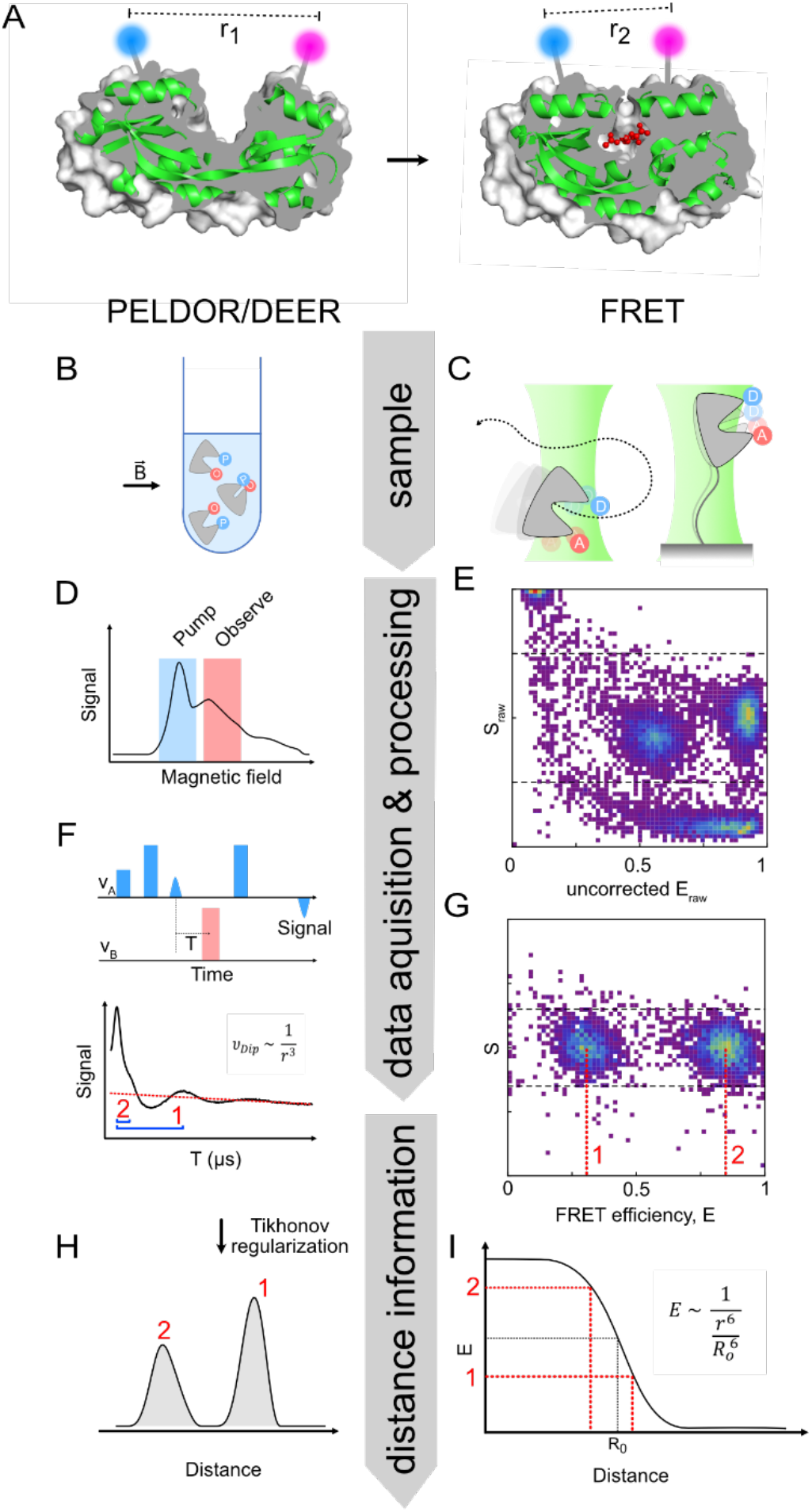
Following conformational changes of proteins via PELDOR/DEER or smFRET. Experimental approaches: **A)** The two methods rely on the presence of either spin centers or fluorescence labels at suitable positions (left: apo HiSiaP, PDB-ID: 2CEY ^13^, right: holo HiSiaP, PDB-ID: 3B50 ^35^). The blue and magenta spheres represent attached labels, between which the distance is either determined by either PELDOR/DEER or smFRET. **Sample preparation: B)** For PELDOR/DEER the experiment is performed with a frozen solution in an EPR tube inside a static magnetic field. (O) and (P) represent observer and pump spins, which are often the same type. **C)** For single-molecule FRET experiments, the labelled protein can either be studied free in liquid solution or coupled to a surface to enable observations over extended time periods. (D) and (A) stand for the donor and acceptor fluorophores of the FRET pair. Further details are shown in Table 1. **Data acquisition & processing: D)** Field-sweep EPR spectrum of the “sample” in B). **F)** By using two different microwave frequencies, subpopulations of the spin centers in the sample can serve as either pump spins or observer spins for the PELDOR/DEER pulse sequence. Integration of the signal at the end of the pulse sequence (blue triangle) for different times T leads to the PELDOR/DEER time trace. The distance information is contained in the different oscillation frequencies of the time trace (indicated by blue brackets). **E)** Typical result of a solution-based smFRET measurement (as shown in C) with two different FRET-populations. The populations in the left upper corner and at the bottom represent proteins with donor- or acceptor-only labelling, respectively. **G)** Sorting out the donor/acceptor only population and applying correction procedures leads to a device-independent FRET efficiency distribution. **Distance information: H)** Time traces can be converted to distance distributions with Tikhonov regularization (DeerAnalysis ^40^). The reliability of the distributions is strongly dependent on the quality and length of the time trace ^41,42^. **I)** The theoretical relation between distance r and FRET efficiency E and the Förster radius of the respective dye pair are used to calculate mean experimental distances between the probes in the structure.

## Results

### A brief comparison of PELDOR/DEER and smFRET

Both PELDOR/DEER and FRET are suitable to measure distances at the nanometer scale and thereby to detect conformational changes of macromolecules in the (frozen) solution state (Figure 1). PELDOR/DEER is a pulsed EPR (electron paramagnetic resonance) experiment, which measures distances by determining the strength of the dipolar coupling between two or more unpaired electrons (spin centers). While PELDOR/DEER is by far the most used experiment for pulsed EPR distance measurements, other pulse sequences have been developed (DQC, SIFTER, RIDME ^43–45^). The applicability of the different pulse sequences depends on the spin centers that are used in the experiment (see below). FRET-assays rely on the dipole–dipole coupling between two spectrally distinct fluorophores to determine the efficiency of non-radiative energy transfer from the electronically excited donor to the acceptor. The energy transfer efficiency depends on the presence of isoenergetic transitions in both molecules (donor emission and acceptor absorption), their relative orientation and the distance between the fluorophores (Figure 1). More recently, it has become possible to study individual donor-acceptor pairs (termed single-pair or single-molecule FRET), which facilitates the conversion of FRET-efficiency to interprobe distance values. Although the two methods are used for similar applications, both have distinct advantages and disadvantages (See Table 1 for an overview). For a full description of each methods’ theoretical background, the reader is referred to the many detailed theoretical reviews and textbooks (e.g. ^46–49^).

**Table 1.** Common practical questions concerning the applicability of PELDOR/DEER and FRET to structural biology questions.

| Structural biology question | PELDOR/DEER | smFRET |
| --- | --- | --- |
| Which range of distances can be measured? | Normally 15-80 Å, but up to 160 Å with fully deuterated proteins | Normally 30-80 Å; 100-150 Å are possible with multiple acceptor dyes |
| How many types of labels are typically needed? | 1 | 2 |
| Can multimeric (homomeric) proteins be studied? | Yes. Multiple distances can be measured in one experiment, using just one type of label and standard equipment. | Yes, but technically demanding. |
| Amount of sample needed? | For Q-band (standard frequency) measurements: 80 µl of ~ 10-30 µM labelled protein | 100 – 400 µl of 15-100 pM labelled protein |
| Physical state of the sample? | Usually frozen solution (50 K). Measurements in aqueous solution are possible but require specific labels (e. g. trityl) and very large or immobilized macromolecules. Depending on the label, other pulse sequences than PELDOR/DEER might be required. | Liquid solution at room temperature and standard cell conditions (37°C) |
| *In vivo* measurements? | Not a standard experiment, especially not under physiological conditions. But, measurements in manually injected frog oocytes or using paramagnetic unnatural amino acids in *E. coli* have been done<sup>52,92</sup>. | Yes |
| Time resolution | Freeze quenched samples can be measured. The time-resolution depends on the freeze quench equipment. | Down to micro- and nanoseconds |
| Time frame for measurements | Normally several hours per measurement for Q-band | Diffusing molecules: 30-60 minutes<br>Immobilized molecules: minutes to hours |

Because most proteins are diamagnetic and devoid of any suitable fluorophores, PELDOR/DEER and FRET experiments usually require the attachment of spin- or fluorescence labels (Figure 1). This is often accomplished by the site-specific introduction of cysteines. The thiol groups of cysteines can be reacted with linker-functionalized dyes or labels, for example via maleimides or thiosulfate esters (see Figure 2 for some typical examples). If the introduction of cysteines is not an option, it is possible to use alternative labelling approaches such as labelled nanobodies ^50^ or unnatural amino acids. The latter can either be fluorescent or paramagnetic themselves or bear functional groups that can be chemoselectively labelled, for instance by click-chemistry ^51–54^. Although the types of labels used for PELDOR/DEER and FRET are quite different, the requirements for suitable labelling positions in proteins are essentially the same: the residue should be solvent-accessible and of no functional importance.

**Figure 2:**
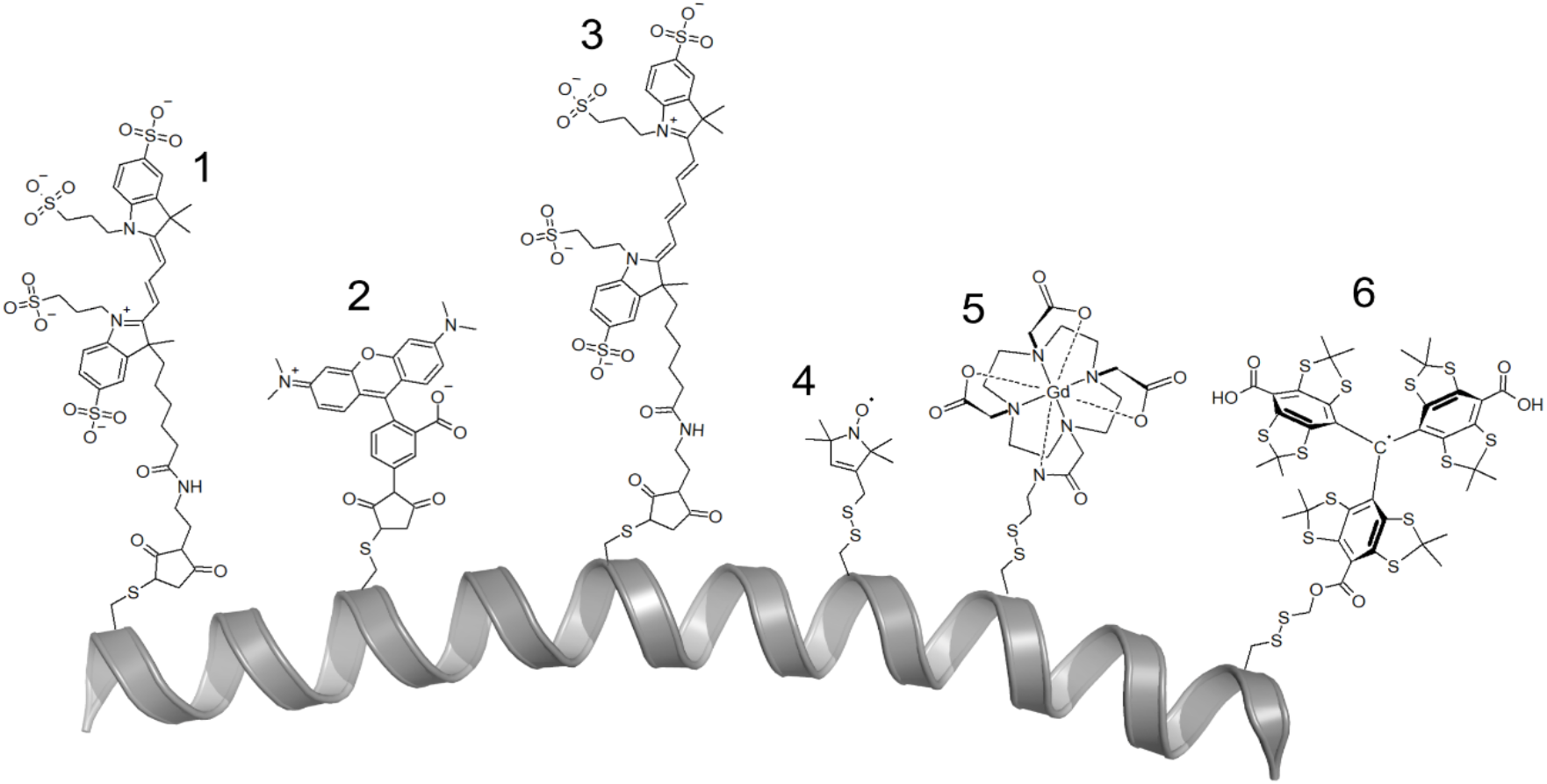
Chemical structures of labels commonly used for smFRET and PELDOR/DEER. **(1)** Maleimide-thiol adducts of Alexa Fluor 555 ^63^ **(2)** TMR (tetramethylrhodamin-5-maleimide) ^64^ and **(3)** Alexa Fluor 647 ^63^ were used in the present study. **(4)** The MTSSL-^65^, **(5)** DOTA-Gd- ^66^ and **(6)** trityl-spin labels ^67^ attached to a cysteine residue via a disulfide bridge. The peptide chain is represented as a grey α helix. Note that many other labels with differing coupling chemistries ranging from click-chemistry to unnatural amino acids have been developed, as well as labels that are specific for nucleic acids.

For PELDOR/DEER spectroscopy the distance between the labels ought to be in the range of 1.5 to 8.0 nm (longer distances of up to 16 nm are accessible with fully deuterated samples) ^46,55,56^. The ideal distance for FRET experiments is around the Förster radius of the particular FRET pair with a typical dynamic range between 3-8 nm (Figure 1F), but in principle, distances up to 10-15 nm can be measured ^57^. Typically, labelling positions are chosen such that the distance change between conformations is as large as possible. In practice, the pool of suitable sites is often surprisingly small. Fortunately, software programs exist to assist in the identification of optimal labelling positions in the case of an available structure or model of the target protein. ^58–62^.

Our aim for the following comparisons was to choose well-established labels and experimental conditions for either method. The PELDOR experiments were thus performed at 50 K with cryo-protected samples using a commercial pulsed Q-band spectrometer. The samples were labelled with MTSSL ^65^. The distance distributions were determined using DeerAnalysis ^40^ and the distance predictions were performed with mtsslWizard ^68^. smFRET experiments of diffusing protein molecules were performed in buffer at room temperature using standard procedures suitable for microsecond alternating laser excition (ALEX) as described before ^63^. The experiments were performed with Alexa Fluor 555 (donor) and Alexa Fluor 647 (acceptor) using a homebuilt confocal microscopy setup with 2-colour detection ^63^. For smFRET assay design, we used a unpublished label analyzer script to identify residues for labelling with optimal dynamic range within the smFRET assay (Gebhardt, Bawidamann, Lipfert & Cordes, unpublished).

### Comparison 1: Sialic acid binding protein HiSiaP

We started with the sialic acid TRAP transporter SBP from *Haemophilus influenzae*. Figure 3A shows a difference distance map of HiSiaP based on the open- and closed crystal structures. The map represents all distance changes between the C-β atoms of the two states.

**Figure 3:**
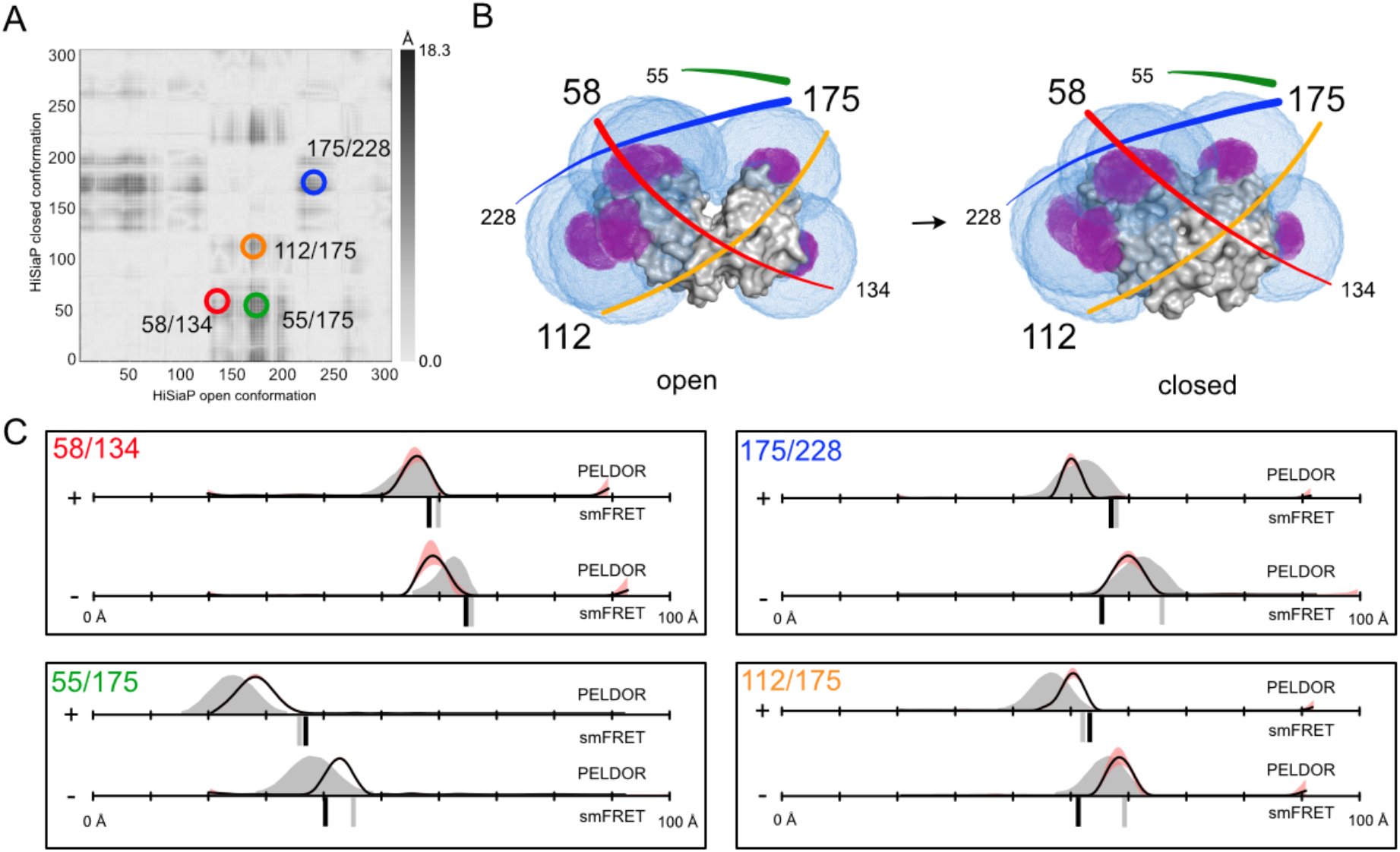
Distance measurements of HiSiaP via EPR and smFRET. **A)** Difference distance map of HiSiaP in open (PDB-ID: 2CEY) and closed (PDB-ID: 3B50) conformation ^69^. The dark spots are protein regions with large conformational changes. The double mutants for distance measurements are highlighted with circles. **B)** Surface presentation of HiSiaP (grey) in open (left) and closed (right) conformation. The accessible volumes of the spin label at six different labelling positions were calculated with mtsslWizard and are represented by magenta meshes. Accessible volumes of FRET label maleimide-Alexa Fluor 647 were calculated with FPS ^70^, and are shown as blue meshes. The double mutants that were used for experiments are illustrated with coloured lines, corresponding to A). **C)** Distance measurements with four different double mutants of HiSiaP without (-) and with (+) substrate. The PELDOR/DEER results are shown above (grey curves for simulation, black curves for experiment) and the FRET distances below the x-axis (grey bars for simulation, black bars for experiment). Raw data for all experiments and confidence interval of PELDOR/DEER distributions are provided in the Supplementary Information. The red shade around the PELDOR/DEER data is the error margin calculated using the validation tool of DeerAnalysis ^40^.

We picked pairs of sites with pronounced distance changes (dark areas) of up to 1.8 nm for labelling. Figure 3B shows the open (PDB-ID: 2CEY) and closed structures (PDB-ID: 3B50) of the protein with the predicted accessible volumes of the spin- (magenta) and FRET- (blue) labels at the selected labelling sites (residues 55, 58, 134, 175, 228). For PELDOR/DEER, all double mutants (58/134, 55/175, 175/228 and 112/175) were labelled with MTSSL, which is by far the most commonly-used spin label for proteins (SI Figure 1). In each case, two PELDOR/DEER measurements were performed, one in the presence (1 mM) and one in the absence of sialic acid (Neu5Ac). Similar to the previously published PELDOR/DEER data for the *Vibrio cholerae* homolog VcSiaP, which shares 49 % amino acid sequence identity, 69 % sequence similarity with HiSiaP ^24^, the EPR-time traces obtained for HiSiaP were of excellent quality with clearly visible oscillations and high signal-to-noise ratios (SI Figure 2). The distance distributions had a single, well-defined peak (Figure 3C, black curves; see SI for distributions with error bars), with a clear shift towards shorter distances in the presence of substrate. The corresponding *in silico* predictions based on the crystal structures (Figure 3C, grey areas), were in good agreement with the experimental PELDOR/DEER-data, considering the known error of ± 3 Å for such predictions ^68^. In summary, for each double mutant, distance changes were measured that were similar in magnitude to those calculated from the crystallographic models, and in agreement with those from the previously published VcSiaP data ^24^.

We next assessed substrate-induced conformational changes in the HiSiaP double mutants by smFRET spectroscopy using Alexa Fluor 555 and Alexa Fluor 647 as donor and acceptor dye, respectively (Figure 2). This FRET pair was chosen for its high photostability, signal intensity and proven compatibility with various protein samples. ^17,28,31^ Labelling quality and sample purity were assessed by size exclusion chromatography, and all samples showed high labelling efficiencies (> 90%) and good donor-to-acceptor labelling ratios (up to ~ 50:50) (SI Figure 3). The experiments were conducted with freely diffusing molecules at ~50 pM concentration to derive mean FRET-efficiency values for the apo- and holo states of HiSiaP. All FRET measurements gave high quality ES-histograms with clearly defined FRET populations (SI Figure 4). However, some of the populations appeared to be broader than expected indicating that they were either composed of molecules with additional conformational flexibility, or there were unwanted photophysical effects arising from the choice of fluorophores and labelling positions (see discussion below).

Figure 3C summarizes the FRET distance measurements in direct comparison with the PELDOR/DEER distance distributions. For smFRET (black bars) only variant 58/134 gave the expected trend for shorter interprobe distances in the presence of ligand in comparison to ligand-free conditions. All other variants (55/175, 175/228, 112/175) failed to reproduce the trends from EPR and structural predictions (grey bars). smFRET data instead suggested that the apo protein adopted a conformation that was more closed than the substrate-bound conformation. Since variant variant 58/134 agreed with both the PELDOR/DEER results and the models based on X-ray structures, it appeared unlikely that a completely unexpected structural feature of the protein was responsible for the observed discrepancies. Considering the known ± 5 Å experimental accuracy of FRET ^71^, one might argue that the two states were simply not discernable for the “offending” double mutants, (contradicting the simulation results in Figure 3C). To examine whether the apo protein was in fact capable of adopting such a “apo-closed” conformation under the conditions in which the FRET experiments were performed, we carried out a burst-variance analysis of three HiSiaP mutants (SI Figure 5). The results showed that the protein exists in a single conformation and is not switching rapidly between distinct states (on the millisecond timescale). These results however do not rule out transitions on a timescale below ~500 μs. The detection of such dynamics would require PIE or MFD analysis of smFRET assays. ^72^ Also, the smFRET experiment gives no information about the possibility that the protein was trapped in a closed state, but data below suggest otherwise. A possible explanation for the discrepancy between crystal structures and smFRET distances for the two label pairs is that the fluorescence labels were partly immobilized by an interaction with a surface feature of the protein. For instance, the sulfonic acid groups of the fluorophores could interact with positively charged patches on the protein surface. Because these effects are highly location dependent, the stochastic labelling combination of donor-acceptor pair and acceptor-donor pair might result in a heterogenous mix of two different types of labelled proteins, i.e., a fluorophore “feels” a different environment depending on the cysteine labelled with it. Interestingly, we could observe such a broadened population ^73^ caused by these two labelling combinations very clearly for a distinct dye-combination (Alexa Fluor 546 – Star 635P) for mutant 112/175 (SI Figure 6). Notably, for all mutants with experimental apo distances that deviated from the simulations, the 1D-E-Histograms showed broadening of the populations indicating artefacts due to fluorophore interaction with the protein surface (SI Figure 4).

To explore the possibility of such unwanted dye-protein interactions, we investigated fluorescence anisotropy and lifetime decays of labelled HiSiaP for two different amino acid positions with a variety of fluorophores. FRET remains a reliable distance ruler for the scenario that at least one of the two fluorophores undergoes free rotation, which is characterized by fast decay of initial anisotropy values and low residual anisotropies. Because the smFRET results for the mutant 58/134 were in good agreement with the simulations, we selected the single mutant at position 58 as a positive control, where we expected low dye-protein interactions. And because all the double mutants containing a fluorophore at position 175 showed unexpected mean FRET values and broad FRET distributions, particularly in the apo state, the single mutant at this position was chosen as a negative example.

The anisotropy decays observed for Alexa Fluor 555 revealed interactions of the dye with the protein at both positions (175 and 58) in both conformational states with residual anisotropies at long delay times just below 0.3 (Figure 4A). Yet Alexa Fluor 555 was fully immobile in the apo conformation for residue 58 (Figure 4A, apo) as seen by the absence of a fast decay component on the sub-nanosecond timescale. Here, we could not identify a short decay component of the time-resolved anisotropy signal on the timescale of fluorophore rotation < 1 ns. For Alexa Fluor 647, both positions showed smaller residual anisotropies (Figure 4B), but also revealed a distinction between apo- and holo state for position 175. In addition, slower anisotropy decays were accompanied by an increase in the fluorescence lifetime of the fluorophores, a finding that was more pronounced for Alexa Fluor 555 ^74^ than for Alexa Fluor 647 (compare apo/holo decays at position 175, Figure 4).

**Figure 4:**
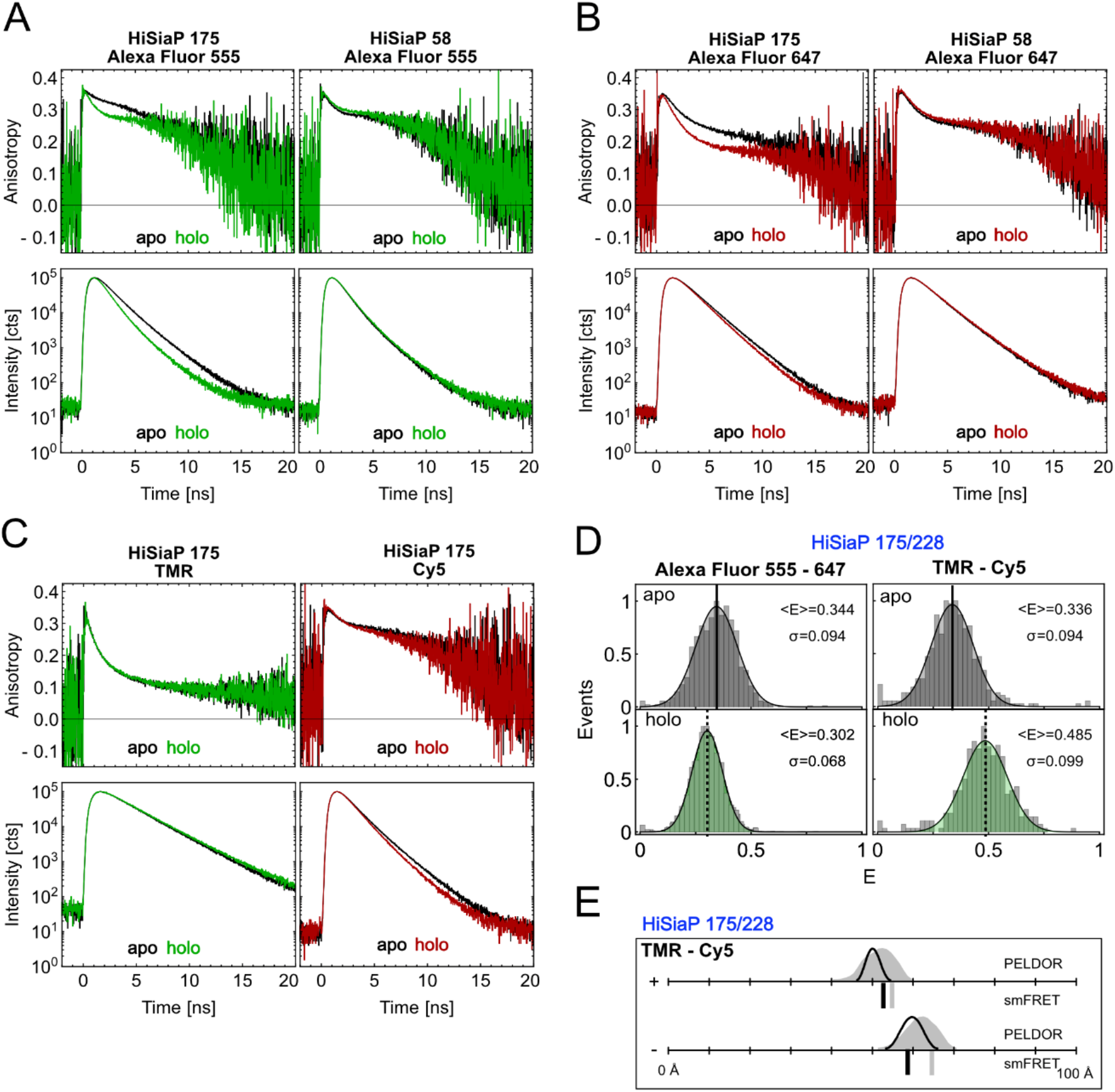
Time resolved fluorescence anisotropy and lifetime measurements on HiSiaP. **A)** Anisotropy decay curves of Alexa Fluor 555 (top row) at residue 175 (left) and 58 (right) and lifetime decay curves (bottom row) under magic angle conditions for apo (black) and holo state (green). **B)** Same measurements as in A) with Alexa Fluor 647 in apo (black) and holo state (red). **C)** Anisotropy decay curves of TMR (top left) and Cy5 (top right) at residue 175 and lifetime decay curves of TMR (bottom left) and Cy5 (bottom right) under magic angle conditions for apo (black) and holo state (colored). **D)** FRET efficiency distributions (center & bottom) of HiSiaP mutant 175/228 for Alexa Fluor 555 – Alexa Fluor 647 (left) and TMR – Cy5 (right) in apo (grey) and holo state (green). **E)** Converted distances from the mean FRET efficiencies are shown as black bars in comparison to simulation (grey bar) and PELDOR/DEER results from Figure 3.

The described photophysical effects with fluorophores at position 175 can explain two observations in our measurements: At first, the larger lifetime increases of Alexa Fluor 555 compared to Alexa Fluor 647 would lead to a broadening of the FRET distribution, because having either the donor or the acceptor at position 175 would result in two distinct FRET states. Secondly, the changes in lifetime (quantum yield), orientation and fluorophore disposition experimentally change the Förster radius and thus impact the proper conversion of FRET-efficiency to distance. In an attempt to avoid these problems, we altered the FRET fluorophore pair to TMR and Cy5 and repeated the experiments for variant 175/228. In contrast to Alexa Fluor 555 and Alexa Fluor 647, these fluorophores are not negatively charged and the linker of TMR is significantly shorter as compared to Alexa Fluor 555 (Figure 2).

The TMR/Cy5 label pair was not our first choice, because it is inferior to Alexa Fluor 555/647 in terms of signal intensity and photostability. Also, for many proteins, charged labels are known to be less prone to stick to the protein surface than hydrophobic labels. In anisotropy and lifetime measurements on the 175 variant in apo and holo state (Figure 4C), however, TMR showed almost ideal behavior with high rotational freedom and an unaffected fluorescence lifetime (Figure 4C), whereas there was a small slowing of the anisotropy decay and increase in lifetime with Cy5 in the apo state. In smFRET experiments using 175/228 with TMR/Cy5, this reduced fluorophore-protein interaction translated into a clear increase in the experimentally determined distance for the apo protein, although this value was still ~5 Å smaller than the simulated distance (Figure 4 D/E). In the holo protein, both the experimental and simulated distance values were decreased by ~4 Å. Taken together, this meant that with the TMR/Cy5 fluorophore pair, the expected substrate-induced domain closure was now observed (Figure 4 D/E), and the absolute distance measurements were in much better agreement with the crystal structures. Thus, it appears that the unexpected FRET results with protein variants labelled at position 175 and the discrepancy between the experimentally determined and simulated distance for the apo protein, were indeed due to interactions of both donor and acceptor fluorophores with the protein surface at this position. It should be noted that also for spin labels, intricate protein/label interactions are known to occur and can explain puzzling results ^75^.

### Comparison 2: Maltose binding protein MalE

MalE has previously been studied by smFRET using Alexa Fluor 555 and Alexa Fluor 647 to elucidate mechanistic implications of conformational dynamics for transporter function (SI Figure 7 and 8) ^31^. In the present work, we used four MalE cysteine double variants (87/127, 36/352, 29/352 and 134/186; Figure 5C and SI Figure 1) with distinct expected distance changes that occur upon ligand binding (see difference distance matrix in Figure 5A). Variants 36/352, 29/352 were designed to show a decrease of distance upon maltose addition. For the 87/127 mutant, the labels were located on the opposite surface of the protein to the substrate binding site near the hinge region, and therefore the expectation from the crystal structures was that the distance would increase for the holo state compared to the apo state. We also included a negative control (134/186), in which the two labels were located in the same domain of the protein (Figure 5C) with the expectation that no substrate-induced distance change occurs (Figure 5C). All variants showed very good agreement between experimental and simulated values in smFRET experiments. Ligand binding was confirmed using microscale thermophoresis as shown previously ^63^.

**Figure 5:**
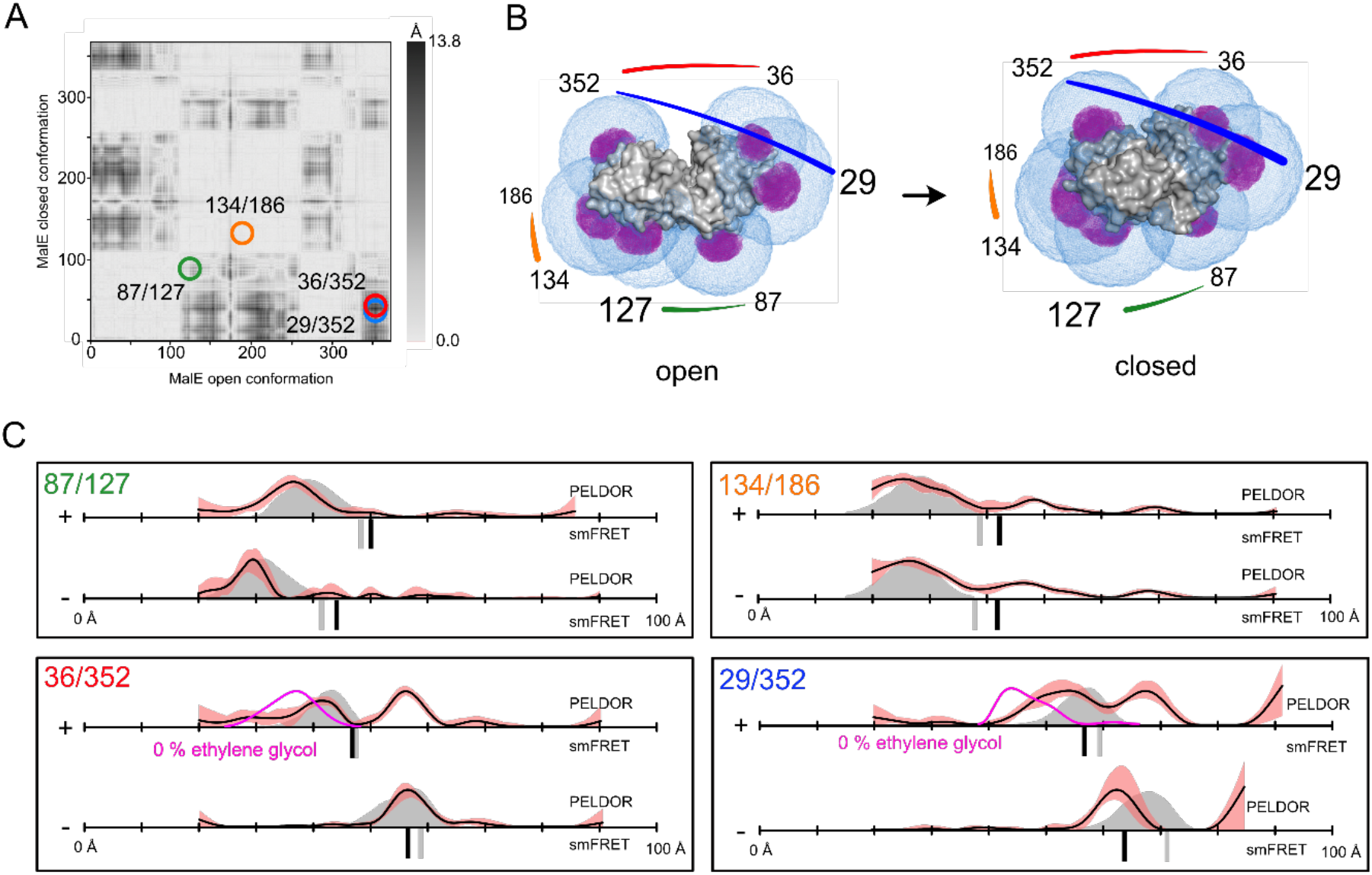
Distance measurements on MalE. **A)** Difference distance map of MalE in open (PDB-ID: 1OMP) and closed (PDB-ID: 1ANF) conformation ^69^. Protein regions with high conformational changes are indicated as dark spots. The double mutants for distance measurements are marked with circles. **B)** Surface presentation of MalE (grey) in open (left) and closed (right) conformation. The accessible volume of the spin label on seven different labelling positions, calculated with mtsslWizard, is represented by magenta meshes. The accessible volume of FRET label maleimide-Alexa647, calculated with FPS ^70^ is shown as blue meshes. **C)** Distance measurements with four different double mutants of MalE without (-) and with (+) substrate. The PELDOR/DEER results are shown above (grey curves for simulation, black curves for experiment) and the FRET distance below the axis (grey bars for simulation, black bars for experiment). PELDOR/DEER results without cryoprotectant are shown as magenta curves. The red shade around the PELDOR/DEER data is the error margin calculated using the validation tool of DeerAnalysis ^40^.

PELDOR/DEER distance measurements using the MTSSL were performed with the same set of mutants (Figure 5). For three of the four MalE variants, this yielded good quality time traces, while the quality for 87/127 was not as high and had relatively low modulation depth and signal to noise ratio (SI Figure 9). For all mutants, except 29/352, the measured apo distances closely matched the predictions obtained from the crystal structure (Figure 5C). Notably, the latter mutant also had the worst match between the simulation and experiment for the smFRET experiments. The addition of 1 mM maltose (Kd for MalE is 1 μM) to the 134/186 mutant protein (our negative control) had, as expected, little effect on the position of the distance peak. Whereas mutant 87/127 seemed to completely close upon substrate addition, the mutant 36/352 revealed what appeared to be a mixture of the holo and apo states. A similar result was obtained for mutant 29/352 with the difference that the “apo distance” in the presence of maltose was ~5 Å longer, and its value was more similar to the simulated apo distance than the distance determined in the actual experiment without maltose. As mentioned above, the qualities of the PELDOR/DEER time traces for the 87/127 mutant were not as high, and the distance change between open and closed conformation was relatively small. Therefore, we cannot exclude the possibility that this mutant also existed in an open-closed mixture after substrate addition (SI Figure 9).

Because binding constants are temperature dependent, and the PELDOR/DEER samples were frozen before the measurement, we checked whether complete closure of MalE was achievable at a higher substrate concentration by repeating the measurements with 10 mM maltose. Within error, these experiments yielded the same mixtures of the holo and apo states as seen with 1 mM maltose (SI Figure 9). Since the lack of complete closure did not result from a sub-saturating maltose concentration in the frozen samples, and it was not observed in the smFRET data, we reasoned that perhaps the cryoprotectant that was used for PELDOR/DEER experiments might be the culprit. Figure 5C shows the PELDOR/DEER results for mutants 36/352 and 29/352 in the presence of 1 mM maltose, and in the presence or absence of 50 % ethylene-glycol cryoprotectant (magenta lines). Further measurements with 25 % ethylene-glycol and glycerol are given in Supplementary Figure 5. When the ethylene glycol concentration was lowered to 25 %, the closed state of MalE already became more prominent and an even stronger change was observed in the absence of cryoprotectant (SI Figure 10). For the measurements without cryoprotectant, the length of the PELDOR/DEER time trace had to be shortened to achieve a good signal to noise ratio (SI Figure 10), and in these cases, a single distance peak with reasonable correspondence to the holo state was observed for both mutants 29/352 and 36/352. Interestingly, the measured distances were shorter than those measured in the presence of cryoprotectant (Figure 5C, red traces).

In summary, for MalE, both methods were able to detect the substrate-induced closure of the protein. A reasonable consistency between the two methods and also in relation to structure-based predictions was found. Initial discrepancies with the PELDOR/DEER measurements were shown to be due to the presence of cryoprotectant.

### Comparison 3: Glutamate/Glutamine binding protein SBD2

Previous to this work, we studied the SBD2 domain of the GlnPQ amino acid transporter by smFRET spectroscopy to elucidate its binding mechanism and its involvement in amino-acid transport ^17^. Here, we conducted smFRET experiments to determine accurate FRET efficiencies for mutants 319/392 and 369/451 using Alexa Fluor 555 and Alexa Fluor 647 (SI Figure 11 and 12). The experimental FRET distances were in good agreement with the *in silico* predictions (Figure 6C, SI Figure 13), where both apo and holo proteins showed a single population. In contrast, in the PELDOR/DEER experiments, the apo form of both SBD2 mutants displayed at least two populations of distances (Figure 6C).

**Figure 6:**
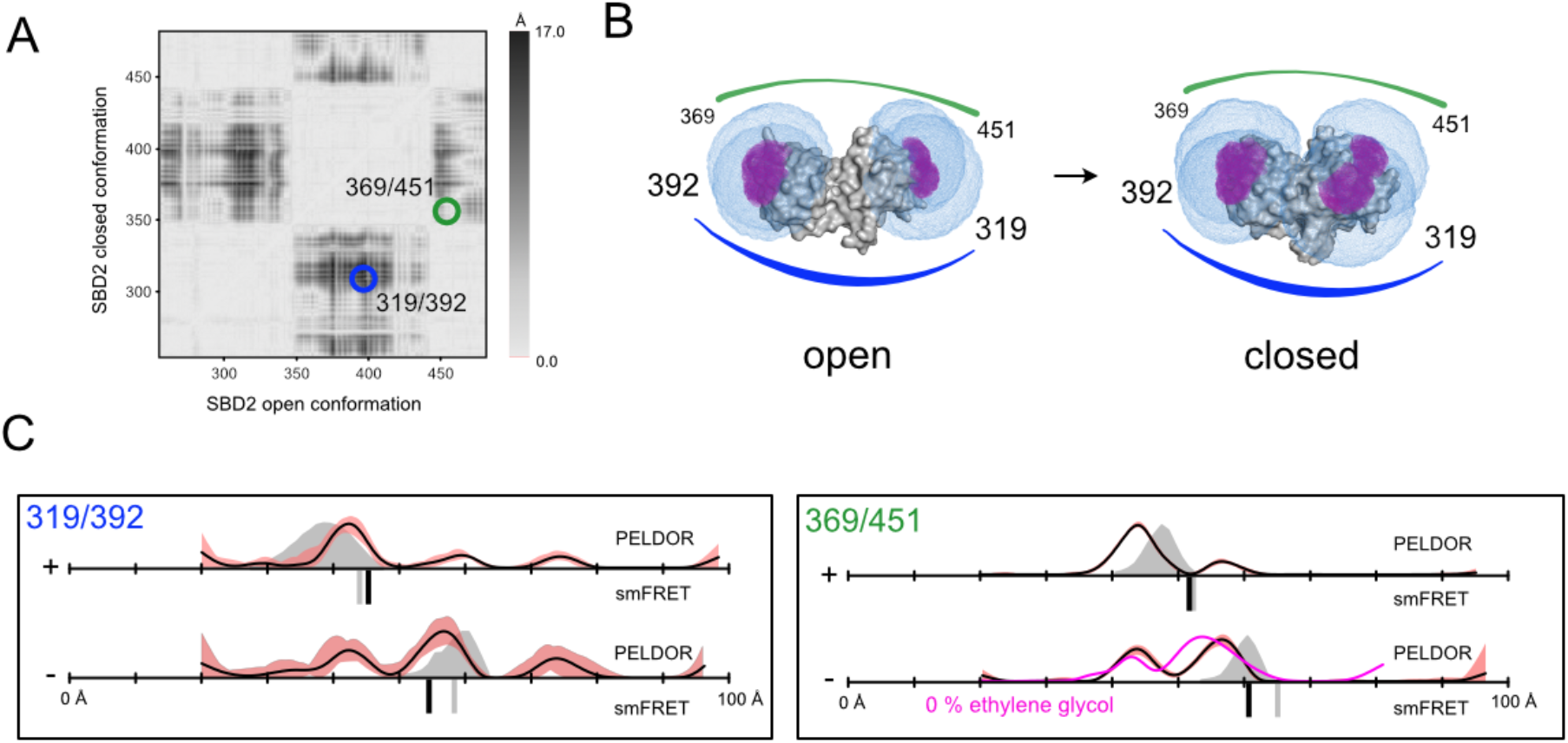
Distance measurements on SBD2. **A)** Difference distance map of SBD2 in open (PDB-ID: 4KR5) and closed (PDB-ID: 4KQP) conformation. Protein regions with high conformational changes are indicated as dark spots. The double mutants for distance measurements are marked with circles. **B)** Surface presentation of SBD2 (grey) in open (left) and closed (right) conformation. The accessible volume of the spin label at four different labelling positions, calculated with mtsslWizard, are represented by magenta meshes and the accessible volume of FRET label maleimide-Alexa647, calculated with FPS ^70^ is shown as blue meshes. **C)** Distance measurements with two different double mutants of SBD2 without (-) and with (+ = 1 mM) substrate. The PELDOR/DEER results are shown above (grey curves for simulation, black curves for experiment) and the FRET distance below the axis (grey bars for simulation, black bars for experiment). PELDOR/DEER results without cryoprotectant are shown as magenta curves. The red shade around the DEER data is the error margin calculated using the validation tool of DeerAnalysis ^40^.

The *in silico* predictions (grey curves) indicated that the major two populations corresponded to the open- and closed conformations. After addition of the substrate glutamine to the protein sample, the PELDOR/DEER distance distributions for both mutants shifted towards shorter distances. However, a small population of ~10 % of the open conformation still appeared to be present (Figure 6C). In this case and within error, the removal of the cryoprotectant during measurements on the apo state of 369/451 had no significant effect on this observation (the time traces are shown in SI Figure 10). To investigate whether the presence of the closed conformation of the apo protein seen with both mutants was due to co-purified glutamine, we performed liquid chromatography mass spectrometry (LC-MS) experiments. Evaluation of the supernatant from purified mutants showed that they did not contain detectable glutamine traces (μM concentrations would be needed to explain our observations, SI Figure 14).

In summary, for the SBD2 protein, both methods were able to discern the open- and closed state. But, the reason for the quantitative differences between the PELDOR/DEER and FRET experiments remained elusive.

### Estimating the influence of linker length on the accuracy of predicted distance distributions

As mentioned above, the discrepancies in our comparisons can be caused by protein-label interactions (see first example HiSiaP), which are extremely challenging to predict. Still, various software program solutions are available that can be used to construct structural models of the labelled proteins and with that to predict inter-label distances ^58–60,76^. Such computer programs predict the accessible volume of a particular label by modelling its structure (or a geometric model thereof) onto the molecular surface of the biomacromolecule. Conformations that are sterically hindered are rejected by the algorithms. Some programs refine the accessible volume by also considering preferred rotameric states of the label ^58,76^ or by favoring states that are close to the molecular surface ^68,70^.

Considering the topic of this work, we asked ourselves, how the prediction uncertainty might be influenced by the different linker length of the typically used spin- and fluorescence labels (Figure 2). To investigate this, we ran a simple simulation with two spherical ensembles, each consisting of 1000 atoms that represent a conformational ensemble of either a spin label or a fluorescence dye (Figure 7A). The radius of the ensemble represents the linker length and its center the position of the Ca atom. We included potential protein-label interactions into this simulation by replacing the position of a particular percentage of the 1000 atoms by the coordinates of one randomly chosen atom. In our model, these cloned atoms represent an immobilized label. We arbitrarily defined a weak interaction to lead to 10 % of the atoms occupying the same position, while the remaining 90 % were randomly distributed, thereby representing an almost freely rotating label.

**Figure 7:**
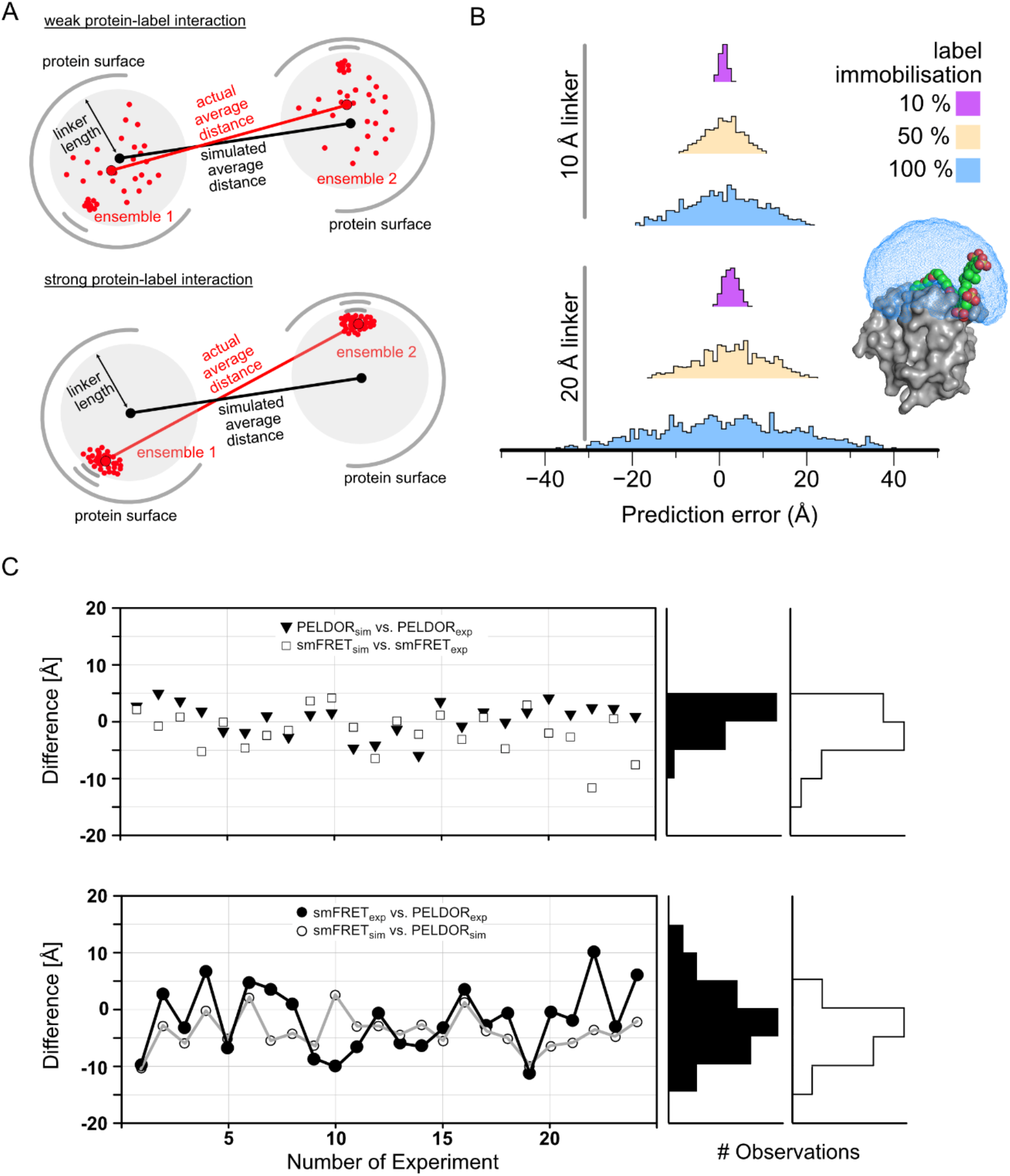
Comparison of PELDOR/DEER and smFRET measurements and the influence of linker length on the correlation between experimental and predicted distances. **A)** Two label ensembles were simulated by distributing 1000 dummy atoms (representing a spin center or fluorophore) in each of two spheres that were located 50 Å apart. The radius of the sphere represents the length of the linker that connects the fluorophore or spin center to the C-alpha atom of the labelled residue. Interactions with the protein surface (grey arcs) are indicated and lead to a clustering of labels at that position. Depending on the degree of interaction between protein and label, the accessible volume approach becomes less accurate. **B)** Histograms of 1000 experiments described in A) with a 10 Å linker (upper plot) and 20 Å linker (lower plot) and varying degree of protein label interaction. The percentage indicates how many percent of the 1000 dummy atoms are localized at the interaction site. As example for a long (20 Å) and immobilized linker, the protein structure of MMP-12 (matrix metalloproteinases, PDB-ID 5L79 ^77^) in conjugation with a Cy5.5 fluorophore (K241, colored spheres) was selected. The surface of the protein is shown in grey and the accessible volume of the fluorophore, calculated with FPS ^70^ is shown as a blue mesh. **C)** The difference between the experimental smFRET and PELDOR/DEER results (black) and the simulated smFRET and PELDOR/DEER results (white).

We then determined the distance between the geometric averages of the 1000 atoms (this corresponds to the experiment) and compared it to the distance of 1000 randomly distributed atoms in the same volume (this corresponds to the accessible volume approach) to calculate the prediction error. This procedure was repeated 1000 times to achieve a statistical distribution of the “interaction site” within the accessible volume. The results are summarized in Figure 7B: For a weakly immobilized label (10 %, magenta), the prediction error was low, even for linkers of 20 Å length (a scenario similar to use of Alexa Fluor labels; Figure 2). This changed markedly when the label interacted more strongly with the protein surface. For instance, if the label occupied a fixed position half of the time, the prediction error increased considerably (50 %, beige). Whereas the error was still acceptable for short linkers, large errors of up to ± 20 Å occurred for longer linkers. If the interaction was even stronger, i.e., for a completely immobilized label (100 %, blue), it was very unlikely to get a useful prediction by the accessible volume approach. An example for such a completely immobilized label is the crystal structure of the matrix metalloproteinases MMP-12 in complex with an fluorophore labelled Cy5.5 inhibitor ^77^ (Figure 7B, PDB-ID: 5L79,), which has a chemical structure related to Alexa Fluor 647 (Figure 2). The label stacks on the surface of the protein and the sulfonic acid groups bind to the positive charges of lysine and arginine residues. Figure 7B shows the size of the accessible volume for the label (blue) for comparison. Note that this interaction is intramolecular and thus not a crystal contact. Similar observations have been made for the R1 nitroxide spin label ^75^.

## Discussion and conclusion

We performed PELDOR/DEER and smFRET experiments on three substrate binding proteins (HiSiaP, MalE and SBD2) to conduct a comprehensive cross-validation of these two important integrative structural biology methods. For this purpose, we used the same labelling sites with both methods and measured the inter-label distances in the presence or absence of their substrates. Both methods showed a good consistency with each other and considering structural models. To get a more quantitative comparison, we took 24 of our measurements (where a single distance peak was observed) and calculated the difference between the average experimental distances. The PELDOR/DEER and smFRET measurements on the same double mutant differed by about 5 Å with an overall spread of ± 10 Å (Figure 7C). To an extent, this difference can surely be addressed to the different label structures and especially to the different linker lengths of the used labels (compare Figure 2). We hence used *in silico* labelling programs, to estimate the extent of these differences ^59,60^. Figure 7C shows that, for most of our measurements, the difference between the two methods is larger (and sometimes much larger) than can be explained by the different linker lengths when a freely rotating label is assumed (otherwise, the black and grey lines in Figure 7C should coincide). Our simulations above (Figure 7A/B) illustrate that even moderate protein label interactions can lead to distance measurements that are seemingly inexplicable (Figure 7) and this is vividly reflected in our first example of HiSiaP using Alexa Fluor 555/647 (Figure 3). Both, experiments and simulations agree that due to the longer linker of the fluorophores, smFRET is likely more prone to this particular problem than PELDOR/DEER (Figure 7).

Looking at Figure 7, three obvious solutions come to mind to improve the situation: (i) Preventing strong interactions between label and protein surface, (ii) the use of shorter linkers and (iii) improving *in silico* labelling approaches. Concerning (ii): Labels with shorter linkers are a topic of constant research. However, for EPR, it is challenging to shorten the length of e.g., the R1 label any further (Figure 1), and short inflexible linkers like the two-armed Rx spin label ^78^ are more likely to disturb the protein structure. Alternative EPR labels such as Gd-, Cu- or trityl with different linker types and lengths are under constant research ^79–81^. For smFRET labels, shorter linkers are not only a problem of chemical synthesis, but also for the implementation of the method itself, because free rotation of the dye is the basic requirement for distance simulations based on accessible volume and exclude orientational effects on FRET efficiency^70^. This requirement would be hard to meet with very short and thus also rigid linkers.

The viability of approach (i) depends on the type of macromolecule. Nucleic acids for instance have a highly negatively charged and predictable surface. So, negatively charged labels (such as many sulphonated dye-molecules) will be repelled and diffuse almost freely in their accessible volume. For proteins, it is much more difficult to predict how the label will interact with the macromolecular surface. It is therefore more straightforward to change the type of label (e.g. charged vs hydrophobic) or to try different labelling positions to validate unexpected or inconsistent results. Indeed, we showed for HiSiaP that switching fluorophores to alter charge and linker length enabled us to correctly detect the expected conformational change.

Approach (iii), an accurate prediction of the label conformations including all solvent molecules and additives by e.g. molecular dynamics simulations would also be a viable solution. However, so far, such time-consuming approaches have not been shown to be any more accurate than the accessible volume approach in large scale benchmarks ^68,82,83^. It should be noted that new and promising approaches to tackle this problem are constantly developed (e.g. ^84 85^). Another option is to measure as many distances as possible and then to investigate, whether a particular observation is truly independent of the labelling site. In this regard, it should also be mentioned that the common practice to re-use the same labelling position for several distance measurements has obvious practical advantages, but renders it much more difficult to validate the different measurements within a single dataset.

Our results on MalE and examples in the literature (e.g. ^86^) demonstrate that the addition of cryo-protectants can influence the result of PELDOR/DEER distance measurements quite significantly. At the high concentrations that are used (10-50% glycerol or ethylene-glycol are typical), the small molecules might interact with the protein and induce a different conformation of the spin label or the protein itself ^86,87^. For smFRET experiments, this is not an issue since the measurements are routinely performed in liquid solution at room temperature. In light of the above, one might argue that the addition of cryo-protectants should simply be avoided. However, without them, it is often not possible to record good quality PELDOR/DEER time traces. The cryoprotectants help to avoid protein aggregation during the freezing procedure and are therefore beneficial for the spectroscopic properties of the PELDOR/DEER sample. They generally lead to increased T2 relaxation times and thereby allow to measure longer distances ^88^. Note, that the cryo-protectant did not significantly influence the distance distributions in previous measurements on a related TRAP transporter SBP with very similar distance results to HiSiaP (Example 1, SI Figure 15), as well as in the SBD2 example or in other studies ^88^. Also, substrate binding proteins such as MalE might be particularly prone to such problems, because they have deep surface crevices that can easily bind small molecules. Nevertheless, especially when unexpected results are found, a control measurement without cryo-protectant or a different cryo-protectant from the large arsenal of such substances should be performed ^89^. Alternatively, the PELDOR/DEER data can be cross validated with a second method such as SAXS ^90,91^. There are constant efforts to develop experimental procedures that allow to reduce the amount of cryo-protectant or to completely avoid their addition, for example by rapid freeze quenching ^86,88^.

Ultimately, it would of course be best to perform PELDOR/DEER measurements at room temperature, since conformational states are likely temperature dependent and therefore the equilibrium of conformational states in the sample might be influenced by the freezing procedure, which, despite all efforts, is still slow compared to the time scale of most molecular motions. This is a possible reason for our observations in MalE and SBD2: The different distances for the closed state in the absence or presence of cryo-protectant might be caused by the different freezing temperature of the respective MalE samples. Similarly, the quantitative differences that were observed in the case of SBD2 might also be caused by the different experimental temperatures. Unfortunately, room temperature PELDOR/DEER measurements (as well as frozen-state FRET measurements) are still challenging experiments and it is therefore no simple task to study the temperature dependence of conformational transitions. However, newly developed labels, such as the trityl spin labels might pave the way for routine room temperature PELDOR experiments ^67^.

Our failure to find the reason for the discrepancies in the SBD2 example is a reminder of the high number of further parameters that can potentially influence the outcome of experiments. One prominent example that was not investigated here is the sample concentration. Whereas smFRET experiments are performed in very dilute solutions, standard Q-band PELDOR/DEER experiments are performed at micromolar concentrations. Considering the molecular crowding in living cells, in-cell measurements would of course be the best option but at the cost of introducing yet more difficult to control parameters into the experiments.

## Conclusion

Both PELDOR/DEER and smFRET are valuable tools in the arsenal of integrative structural biology. While we found overall agreement between the two methods, our experiments also revealed some surprising discrepancies. We found that these are caused by the different experimental approaches, including the nature of the used labels, experimental temperatures and the differences between a single molecule vs bulk point of view. For both techniques, the largest uncertainty stems from the use of labels and the question whether the label undergoes interaction with the protein surface. Due to the longer linkers of FRET labels, this problem is inevitably more prominent for smFRET. For PELDOR/DEER on the other hand, the use of cryoprotectant led to artefacts in one of our examples. Thus, it is important to keep in mind that also the “orthogonal methods” that are used to circumvent the limitations of crystallography and EM are not perfect and that results that appear to falsify high-resolution structures derived from e.g. cryo-EM or X-ray crystallography should be cross-validated and not be taken at face value.

## Author contributions

GH and TC conceived and supervised this study. MFP performed the PELDOR/DEER experiments, JG performed PELDOR/DEER experiments on VcSiaP. CG performed single-molecule FRET experiments. RM and MFP provided protein samples. MFP, CG, GHT, TC and GH have designed the experiments and analyzed the data. MFP, CG, TC and GH wrote the manuscript. All authors contributed to the discussion of the results and the final version of the manuscript.

## Acknowledgments

This project was financed by the German Research Foundation (Deutsche Forschungsgemeinschaft, DFG) in projects no. HA 6805/4-1 and HA 6805/5-1 (to GH), GRK2062 project C03 (to TC), SFB863 project A13 (to TC), the Center for integrated protein science Munich CiPSM (start-up funding to TC), an ERC starting grant ERC-StG 638536 SM-IMPORT (to TC), and the Center of Nanoscience Munich (project funding to TC). MFP acknowledges a PhD fellowship from the Konrad Adenauer Stiftung. CG acknowledges a PhD fellowship from the Studienstiftung des deutschen Volkes. MFP, JG and GH thank Prof. Olav Schiemann, University of Bonn, for support and access to the EPR spectrometers and D. Abdullin for discussions and helpful comments on the manuscript. TC thanks Nicola Gericke for help with purification of substrate binding domain 2. GH and MFP thank Dr. Frank Eggert and Prof. Stefanie Kath-Schorr for help with the MS analysis. We thank E. Lerner for the gift of HP3 and D. Griffith for proofreading and commenting on the manuscript.

## Material & Methods

### Selection of labelling sites

Dependent on the particular method (PELDOR/DEER or smFRET) for which the protein mutants were produced, we used different software to calculate suitable labelling positions. For spin label positions we used mtsslSuite (www.mtsslsuite.isb.ukbonn.de) and calculated a difference distance map between the open and closed conformation (as shown in results) ^69^. With this map we identified regions with large conformational changes and select amino acids inside these regions, which are located on the surface of the protein to obtain a good accessibility. For smFRET studies, optimized double cysteine mutants were created with a yet unpublished software-tool for optimal fluorophore labelling (Gebhardt & Cordes, unpublished), which will be described in a forthcoming publication. In short: residues were rated based on different parameters such as solvent exposure or conservation to obtain a labelling feasibility estimate. Residues with high ratings are paired to find good smFRET pairs with large distance change between apo and holo (or no distance change as negative control).

### Protein expression and purification

The TRAP SBPs HiSiaP and VcSiaP were expressed and purified as described before ^24^. To prevent co-purification of the substrate, the *E. coli* cells were cultured in M9 minimal medium. For purification, the protein was loaded onto a benchtop Ni-affinity chromatography, followed by an ion exchange chromatography and a size exclusion chromatography. In all steps the buffers were supplemented with 1 mM Tris(2-carboxyethyl)phosphine (TCEP) to avoid dimerization of the cysteine mutants and the purity was checked after each step with SDS-PAGE. The purified protein solution was concentrated to 20 mg/mL and stored at −80 °C until labelling. SBD2 and MalE were expressed and purified as described before ^31^.

### LC-MS

The LC-MS analysis was performed on an HTC esquire (Bruker Daltonic) in combination with an Agilent 1100 Series HPLC system (Agilent Technologies). Analysis gradient: 5→100% MeCN (solvent B)/0.1%formic acid (solvent A) in 20 min at a flow rate of 0.4 mL min-1 using a Zorbax Narrow Bore (2.1×50 mm, 5 μm) C18 column (Agilent Technologies).

### Protein labelling

<u>Spin labelling</u>: In the first step, the reducing agents in the protein solution were removed with a PD10 desalting column (GE Healthcare), using a buffer based on 50 mM Tris, 50 mM NaCl, pH 8 without TCEP or DTT. Immediately after elution from the column, the protein eluate was treated with 5 times excess per cysteine of the nitroxide spin label MTSSL (Toronto Research Chemicals, Canada), dissolved in DMSO. The labelling was carried out for one hour at room temperature under gentle shaking. Afterwards, the protein was concentrated and another PD10 desalting column was used to remove free spin label. The protein eluate was again concentrated to ~ 20 mg/mL. The labelling was verified and quantified with continuous-wavelength EPR spectroscopy (cw-EPR) ^93^ on an EMXnano X-band EPR spectrometer from Buker (Billerica, MA). The spin labelled proteins were dilute to a concentration of 25 μM with standard buffer and a total volume of 10 μL sample was prepared into a glass capillary, sealed with superglue. The magnetic field of the cw-EPR spectrometer at room temperature were set to a center field of 3448 G and the microwave frequency to 9.631694 GHz. The microwave power was set to 2.5 mW, the power attenuation to 16 dB and the receiver gain to 68 dB. The cw-EPR spectra were recorded with a sweep width of 150 G, a sweep time of 10.03 s with 20.48 ms time constant and 1 G modulation amplitude. For every sample 350 cw-EPR spectra were averaged to obtain a good signal to noise ratio. The concentration of the spin label and the labelling efficiency were determined with the Bruker software Xenon by double integration of the cw-EPR spectrum.

<u>Fluorophore labelling</u>: Proteins were labelled as described previously ^17,28^. The cysteines were stochastically labelled with the maleimide derivative of the dyes TMR, Alexa Fluor 555, Alexa Fluor 647 and Cy5 (ThermoFischer Scientific). His-tagged proteins were incubated in 1 mM DTT to keep all cysteine residues in a reduced state and subsequently immobilized on a Ni Sepharose 6 Fast Flow resin (GE Healthcare). The resin was incubated 2-4 h at 4°C with 25 nmol of each fluorophore dissolved in 1 ml of labelling buffer 1 (50 mM Tris-HCl pH 7.4, 50 mM KCl, 5% glycerol) and subsequently washed sequentially with 3 ml labelling buffer 1 and buffer 2 (50 mM Tris-HCl pH 7.4, 150 mM KCl, 50 % glycerol) to remove unbound fluorophores. Bound proteins were eluted with 500 ml of elution buffer (50 mM Tris-HCl pH 8, 50 mM KCl, 5% glycerol, 500 mM imidazole) The labelled protein was further purified by size-exclusion chromatography (ÄKTA pure, Superdex 75 Increase 10/300 GL, GE Healthcare) to eliminate remaining fluorophores and remove soluble aggregates. For all proteins, labelling efficiencies were higher than 70% and donor-acceptor pairing at least 20%.

### PELDOR/DEER spectroscopy

If not indicated in the results, the standard EPR samples were prepared and the measurements were set up as described in the following. The proteins and additives were mixed and diluted to a concentration of 15 μM in a volume of 40 μL with PELDOR/DEER buffer (100 mM TES pH 7.5, 100 mM NaCl in D2O). The substrate concentrations were chosen to 1 mM *N*-acetyl neuraminic acid for HiSiaP, 1 mM maltose for MalE and 100 μM glutamine for SBD2. The solutions were supplied with 40 μL deuterated ethylene glycol, transferred into a 3 mm quartz Q-band EPR tube and immediately flash-frozen and stored in liquid nitrogen.

The PELDOR/DEER experiments were measured on an ELEXSYS E580 pulsed spectrometer from Bruker in combination with an ER 5106QT-2 Q-band resonator. The temperature was set to 50 K with a continuous flow helium cryostat (CF935, Oxford Instruments) and a temperature control system (ITC 502, Oxford Instruments). The PELDOR/DEER time traces were recorded with the pulse sequence π/2(υ_A_)-τ_1_-π(υ_A_) − (τ_1_+t) − π(υ_B_)-(τ_2_-t)-π(υ_A_)-τ_2_-echo. The frequency υ_A_ of the detection pulses were set 80 MHz lower than the frequency of the pump pulse υ_B_, which was set to the resonator frequency and the maximum of the nitroxide spectrum. Typically, the short repetition time was 1000 μs and the lengths of τ_1_ and τ_2_ was 12 and 24 ns, respectively. The contribution of deuterium ESEEM to the PELDOR/DEER time trace was suppressed by addition of 8 observed time traces with variable τ_1_ time (Δ = 16 ns). The PELDOR/DEER background was fitted by a monoexponential decay. The distance distributions were calculated and validated by means of DeerAnalysis 2018 ^40^.

### smFRET spectroscopy

Solution based smFRET experiments were performed on a homebuilt confocal ALEX microscope as described in ^94^. All sample solutions were measured with 100 μl drop on a coverslip with concentration of around 50 pM in buffer 1. The fluorescent donor molecules are excited by a diode laser OBIS 532-100-LS (Coherent, USA) at 532 nm operated at 60 μW at the sample in alternation mode (100 μs alternation period). The fluorescent acceptor molecules are excited by a diode laser OBIS 640-100-LX (Coherent, USA) at 640 nm operated at 25 μW at the sample. The lasers are combined and coupled into a polarization maintaining single-mode fiber P3-488PM-FC-2 (Thorlabs, USA). The laser light is guided into the epi-illuminated confocal microscope (Olympus IX71, Hamburg, Germany) by dual-edge beamsplitter ZT532/640rpc (Chroma/AHF) focused by a water immersion objective (UPlanSApo 60x/1.2w, Olympus Hamburg, Germany). The emitted fluorescence is collected through the objective and spatially filtered using a pinhole with 50 μm diameter and spectrally split into donor and acceptor channel by a single-edge dichroic mirror H643 LPXR (AHF). Fluorescence emission was filtered (donor: BrightLine HC 582/75 (Semrock/AHF), acceptor: Longpass 647 LP Edge Basic (Semroch/AHF), focused on avalanche photodiodes (SPCM-AQRH-64, Excelitas). The detector outputs were recorded by a NI-Card PCI-6602 (National Instruments, USA).

Data analysis was performed using home written software package as described in ^17^. Single-molecule events were identified using an All-Photon-Burst-Search algorithm with a threshold of 15, a time window of 500 μs and a minimum total photon number of 150 ^95^. Photon count data were extracted, background subtracted, corrected for spectral crosstalk, and different quantum yields / detection efficiencies described in ^71,96^. An exemplary correction procedure of all correction steps is illustrated in SI Figure 16.

E-histogram of double-labelled FRET species with Alexa Fluor 555 and Alexa Fluor 647 was extracted by selecting 0.25<S<0.75. E-histograms of open state without ligand (apo) and closed state with saturation of the ligand (holo) were fitted with a Gaussian distribution 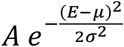. The burst variance analysis (BVA) ^97^ was performed on the same data with a photon binning of 5 photons for selected bursts with stoichiometry 0.25<S<0.75.

### In silico distance simulations

For both methods, we used available programs and combined the *in silico* distance simulations with the experimental distances in the result parts. For PELDOR/DEER simulations, we used mtsslWizard (www.mtsslsuite.isb.ukbonn.de) where an ensemble of rotamers is calculated for each labelling position by rotation of the bonds from the spin label (see results) ^68^. After this, the average distance and the distance distribution between two of these ensembles can be determined.

For smFRET we used the FRET-restrained positioning and screening method established by the Seidel lab ^60^. This method allows the determination of a FRET-efficiency-averaged model distance between the two dyes using the crystal structure information. For distance simulations we employed a simple dye model, in which three parameters were used to determine the accessible volume the dye can sample: (i) linker-length (linker), linker-width (W), and the fluorophore volume, which can be derived from an ellipsoid using R1, R2 and R3. With this information, the average distance between two of these spheres was calculated. The dye parameter for the different fluorophores are shown in Table 2. An average distance was calculated with the FPS software by exchanging donor and acceptor positions and vary the linker length (±1 Å) as well as linker width and radii (±0.5 Å).

**Table 2:** Geometric Parameters for in silico predictions of FRET labels.

| Label | Linker linker [Å] | W [Å] | R1 [Å] | R2 [Å] | R3 [Å] |
| --- | --- | --- | --- | --- | --- |
| **Alexa Fluor 555 – C2 Maleimide** <sup>63</sup> | 21 | 4.5 | 8.8 | 4.2 | 1.5 |
| **Alexa Fluor 647 – C2 Maleimide** <sup>60</sup> | 21 | 4.5 | 11 | 4.7 | 1.5 |
| **TMR** <sup>60</sup> | 12 | 4.5 | 6 | 4.2 | 1.5 |
| **Cy5** <sup>60</sup> | 21 | 4.5 | 11 | 3 | 1.5 |

### Fluorophore lifetime and time-resolved anisotropy measurements

Lifetime and anisotropy decay measurements were performed as described in ref. ^98^ (ch. 11) on a homebuilt setup ^63^: 400 μl of sample is measured in a 1.5×10 mm cuvette at a concentration of around 100 nM. The sample is excited by a pulsed laser (LDH-P-FA-530B for green fluorophores/LDH-D-C-640 for red fluorophores with PDL 828 “Sepia II” controller, Picoquant). Excitation polarization is set with a lambda-half-waveplate (ACWP-450-650-10-2-R12 AR/AR, Laser Components) and a linear polarizer (glass polarizer #54-926, Edmund Optics). Emission light is polarization filtered (wire grid polarizer #34-315, Edmund Optics). The emission light is collected with a lens (AC254-100-A, Thorlabs) and scattering light is blocked with filters (green: 532 LP Edge Basic & 596/83 BrightLine HC, AHF; red: 635 LP Edge Basic & 685/80 ET Bandpass, AHF). The signal is recorded with an avalanche-photodiode (SPCM-AQRH-34, Excelitas) and a TCSPC module (HydraHarp400, Picoquant). Polarization optics is mounted in homebuilt, 3D-printed rotation mounts and APD is protected from scattered light with a 3D-printed shutter unit.

The excitation power was 10 μW and the concentration was finetuned to have ~50 kHz count rate under magic angle conditions. All anisotropy and lifetime measurements were recorded for 5 min in the order vertical (VV1), horizontal (VH1), magic angle (MA), horizontal (VH2), and vertical (VV2) under vertical excitation. Anisotropy was calculated based on the two vertical and horizontal measurements by taking the mean values to compensate for small drifts in laser power or slow changes in fluorophore concentration due to sticking. With 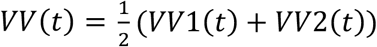 and 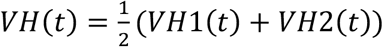, we obtain the anisotropy decay as 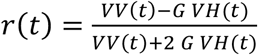.

## Supporting Information

**SI Figure 1:**
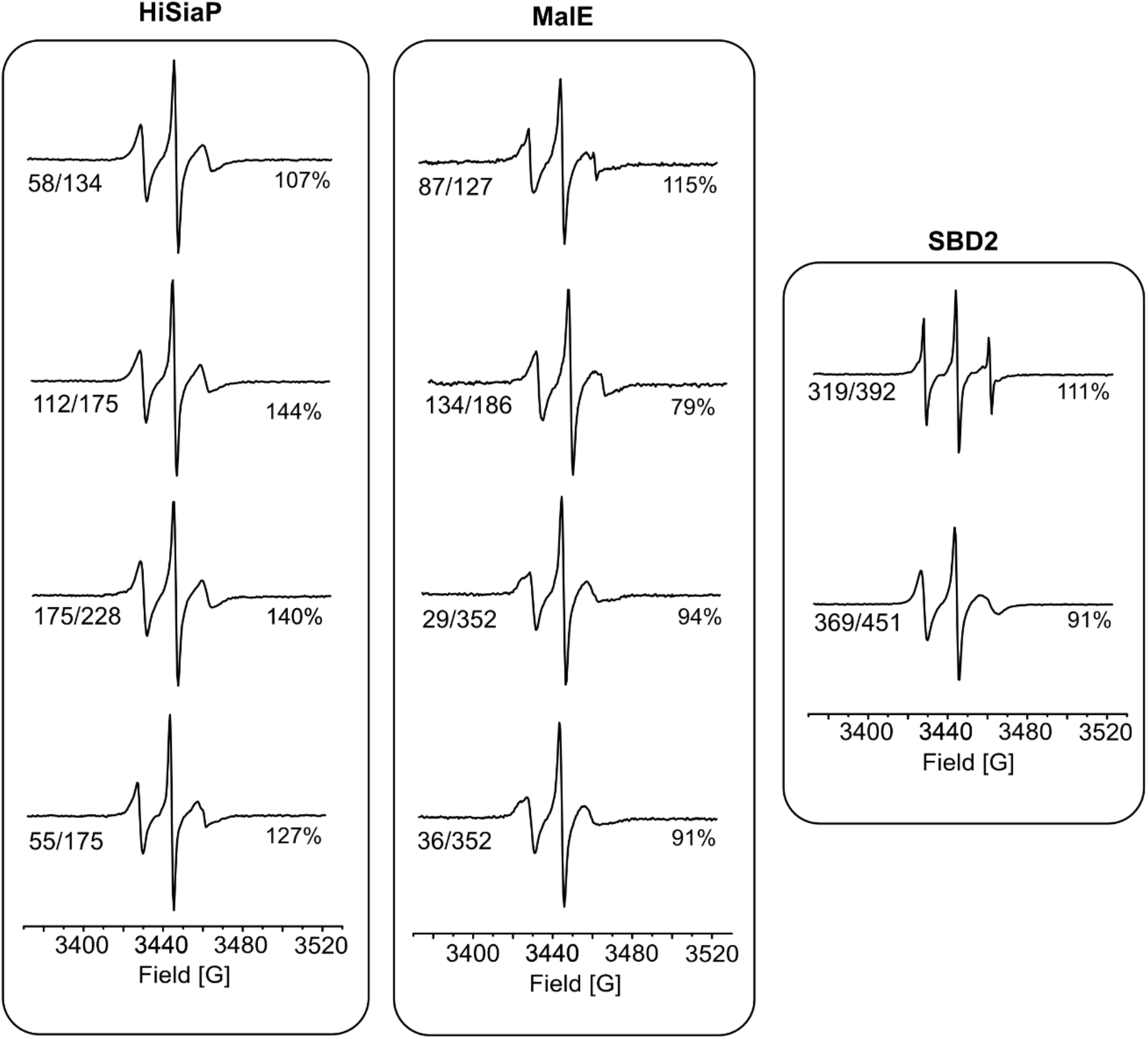
cw-EPR spectra of spin labelled double mutants. The labelling of each mutant with MTSSL was verified with room temperature cw-EPR spectroscopy (X-band). The labelling efficiencies were determined with the spectrometer software and is given next to the spectra.

**SI Figure 2:**
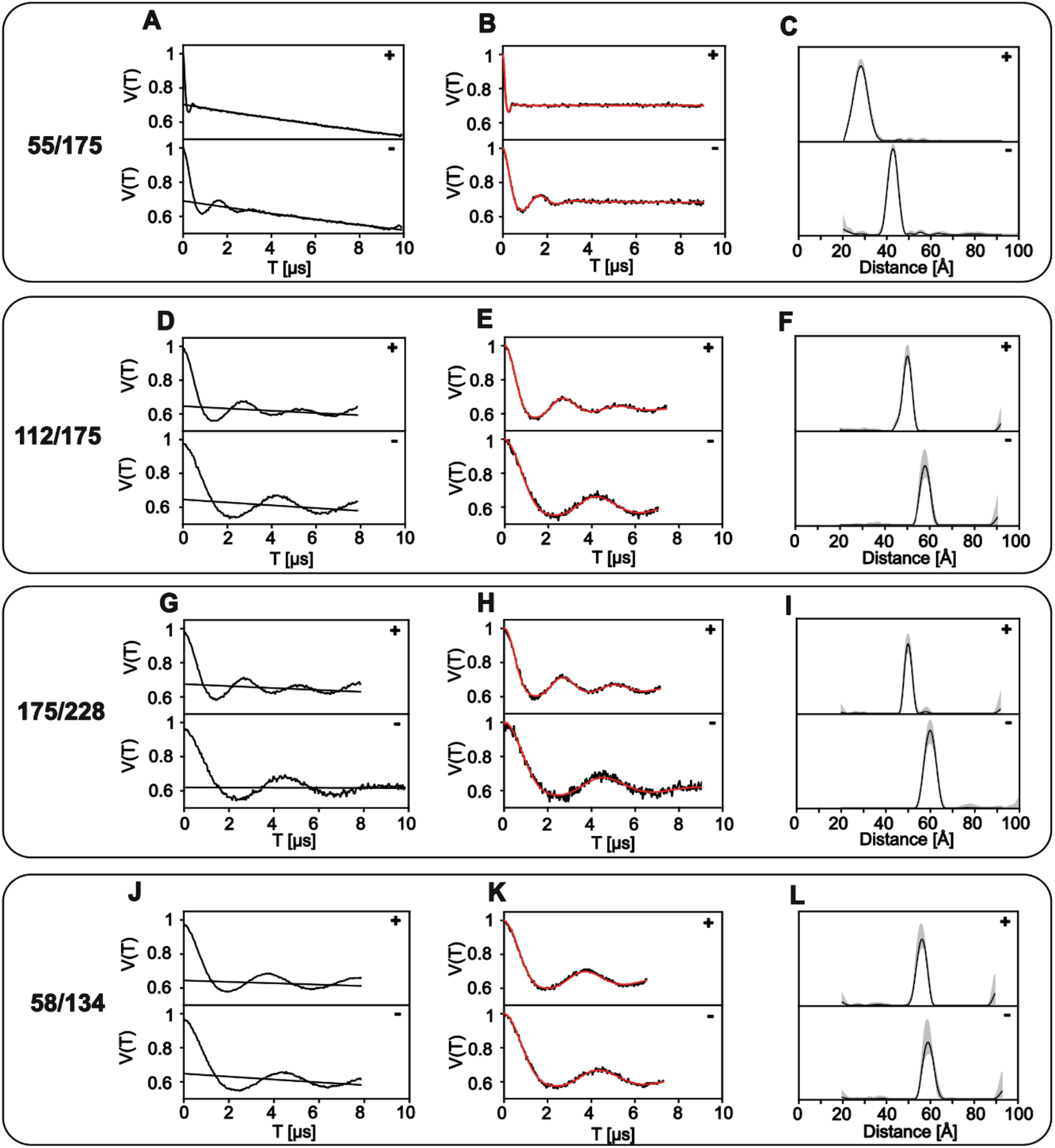
PELDOR/DEER data of HiSiaP mutants. **A, D, G, J)** Raw PELDOR/DEER time traces for apo (-) and holo (+) measurements of each double mutant. The background, which was used for correction of the signal, is indicated as a black line. **B, E, H, K)** Background-corrected PELDOR/DEER time traces (black) and fits of the signal (red). **C, F, I, L)** Distance distributions from PELDOR/DEER time traces (black) with validation of the distribution (grey).

**SI Figure 3:**
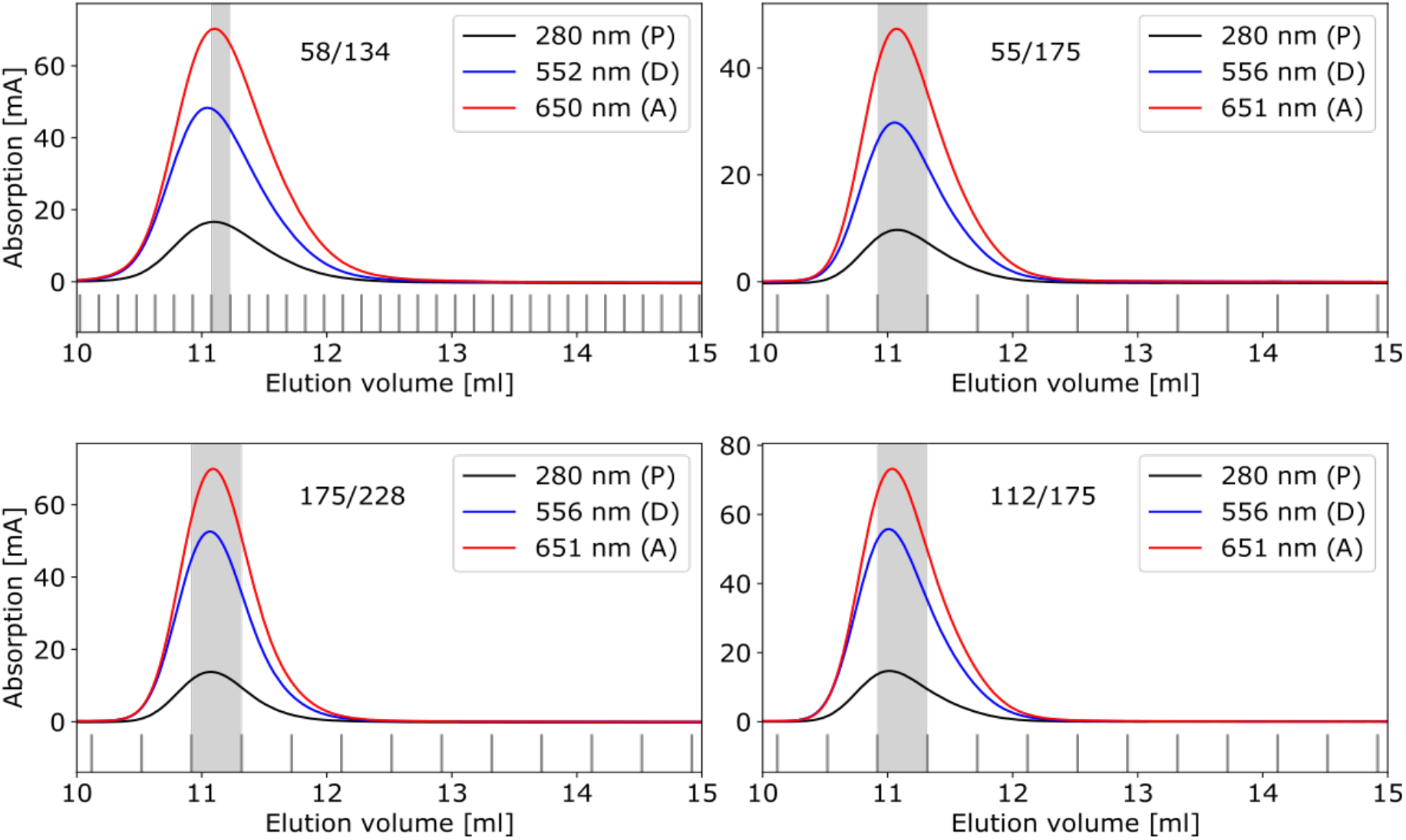
Size exclusion chromatography of HiSiaP mutants labelled with Alexa Fluor 555 – Alexa Fluor 647. Absorption profile of the size extrusion chromatography (ÄKTA, Superdex 75 Increase 10/300 GL, GE Healthcare) for all tested HiSiaP mutants 58/134, 175/228, 55/175 and 112/175 to monitor protein concentration (280 nm) and Alexa Fluor 555 (552 nm) / Alexa Fluor 647 (650 nm). The grey area indicates the fraction used in the smFRET experiments, where labelling efficiencies of >90% was achieved for all samples.

**SI Figure 4:**
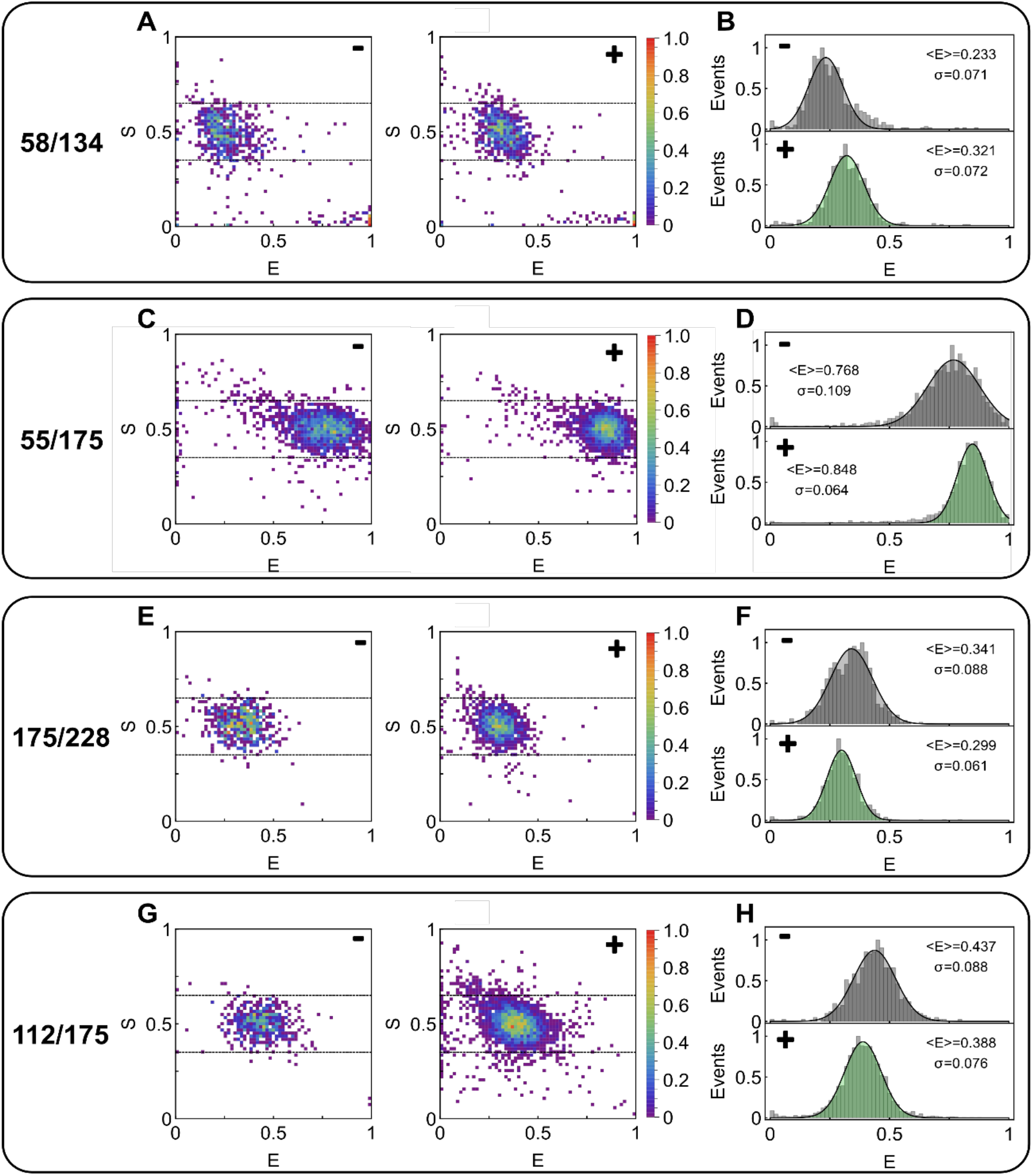
smFRET data of HiSiaP with Alexa Fluor 555 – Alexa Fluor 647. **A, C, E, G)** ES-2D-Histograms of HiSiaP mutants 58/134, 175/228, 30/175, 55/175, and 112/175 in apo state (-) and holo state (+). **B, D, F, H)** 1D-E-Histograms extracted from the ES-data for apo (grey) and holo (green) are fitted with a 1D-Gaussian distribution. Mean <E> and standard deviation σ are labelled.

**SI Figure 5:**
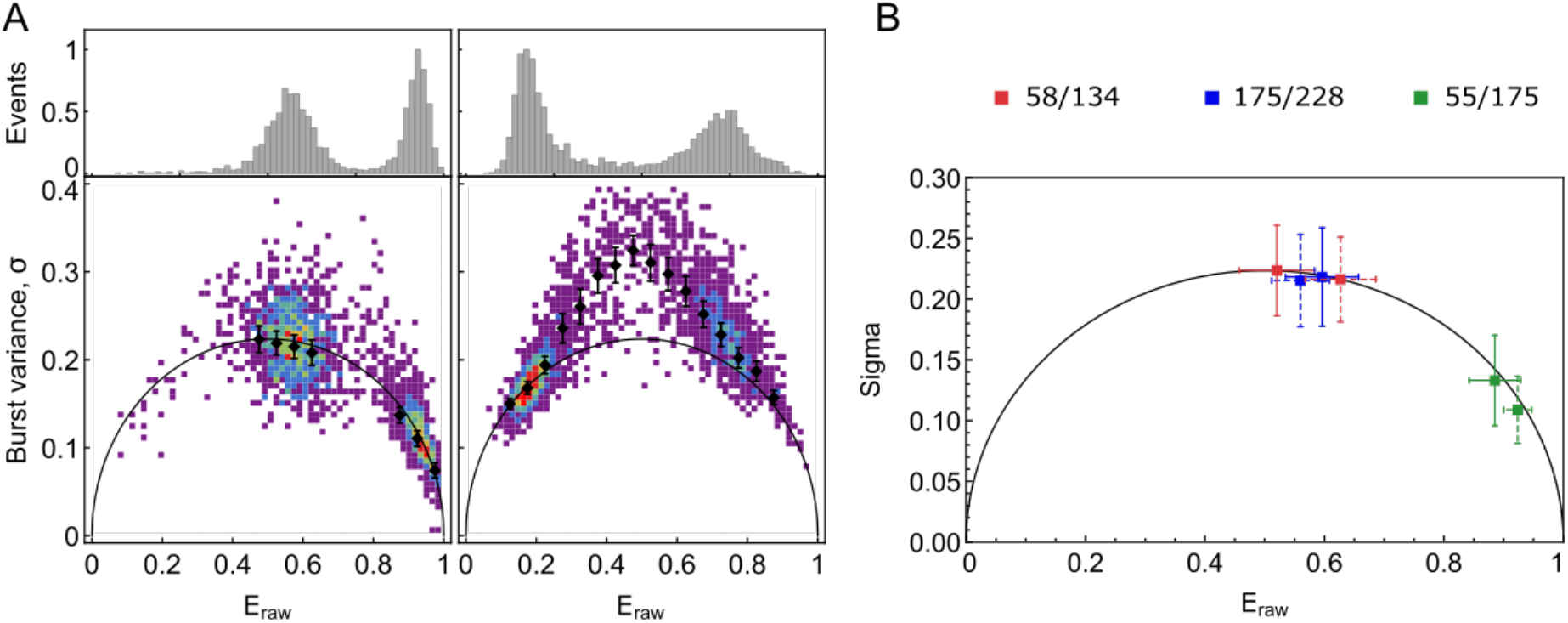
Burst-variance analysis of HiSiaP ALEX data. **A)** Burst variance analysis example of 55/175 and 175/228 in their holo states (left data set) and dynamic control experiments with a fluctuating DNA-hairpin (right data set) from ref. ^99^. Data are binned into bins of 0.05 and mean and standard error of mean are shown (black). **B)** Population mean and standard deviation of all bursts of one measurement of burst variance analysis for three HiSiaP mutants from Figure 3 in apo (solid) and holo state (dashed).

**SI Figure 6:**
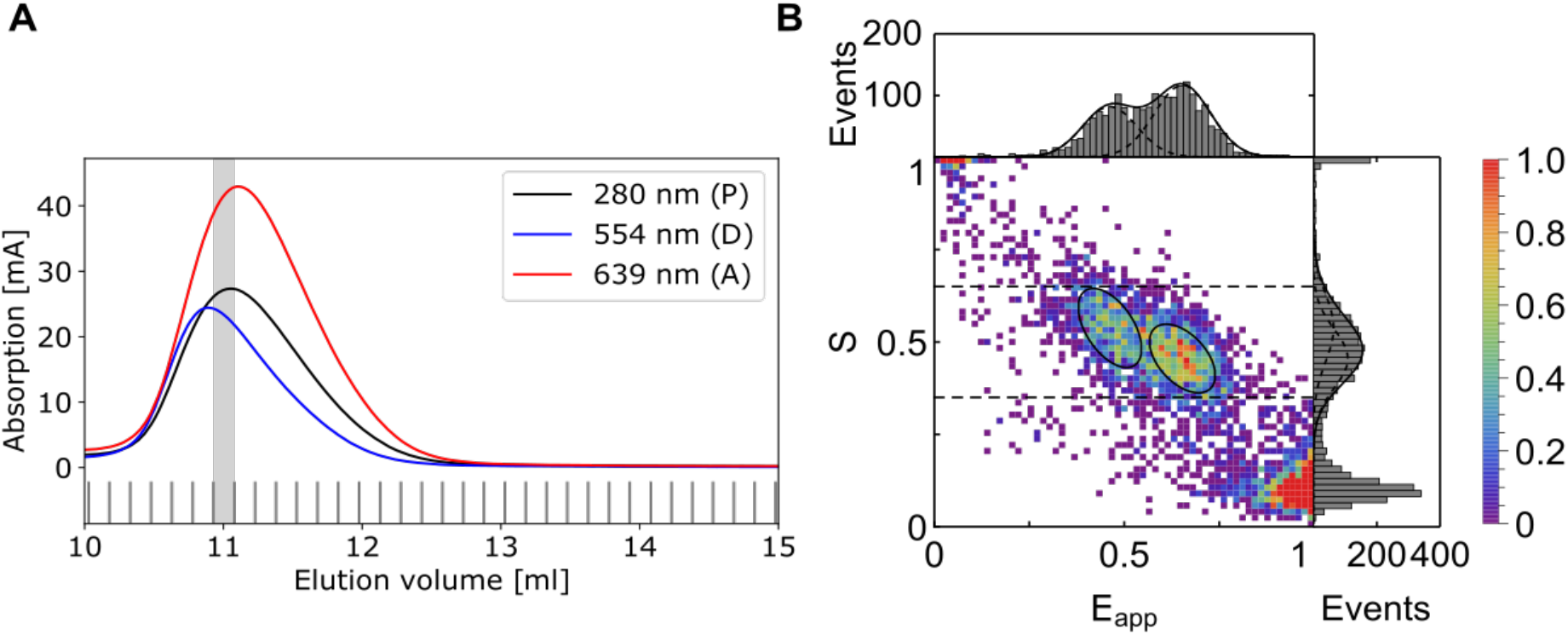
Extreme example for environmental effect on FRET efficiency. HiSiaP mutant 112/175 labelled with Alexa Fluor546 – Star635P. **A)** Absorption profile of the size extrusion chromatography (ÄKTA, Superdex 75 Increase 10/300 GL, GE Healthcare) to monitor protein concentration (280 nm) and Alexa Fluor546 (554 nm) / Alexa Fluor647 (639 nm). The grey area indicates the fraction used in the smFRET experiments, where labelling efficiencies of >90%. **B)** ES-2D-Histograms of mutant from A) in apo state. 1D-E-Histograms and 1D-S-Histogram are shown on top and on the right, respectively. The ES-data are fitted with a 2D-Gaussian distribution where the 1D-integrals are shown in the 1D-histograms (black lines).

**SI Figure 7:**
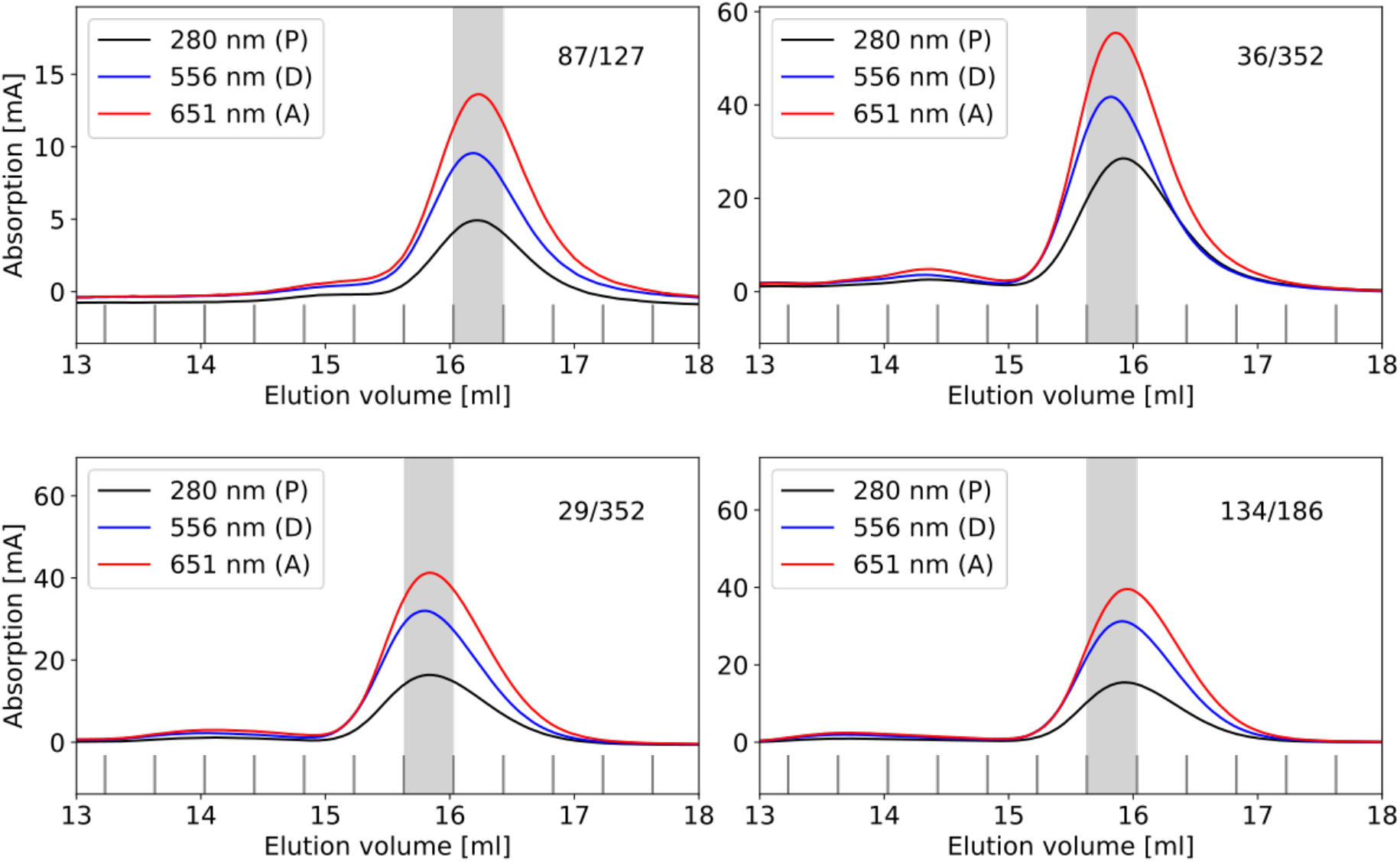
Size exclusion chromatography of MalE mutants labelled with Alexa Fluor 555 – Alexa Fluor 647. Absorption profile of the size extrusion chromatography (ÄKTA, Superdex 75 Increase 10/300 GL, GE Healthcare) for all tested MalE mutants 87/127, 36/352, 29/352 and 134/186 to monitor protein concentration (280 nm) and Alexa Fluor 555 (552 nm) / Alexa Fluor 647 (650 nm). The grey area indicates the fraction used in the smFRET experiments, where labelling efficiencies of >90% was achieved for all samples.

**SI Figure 8:**
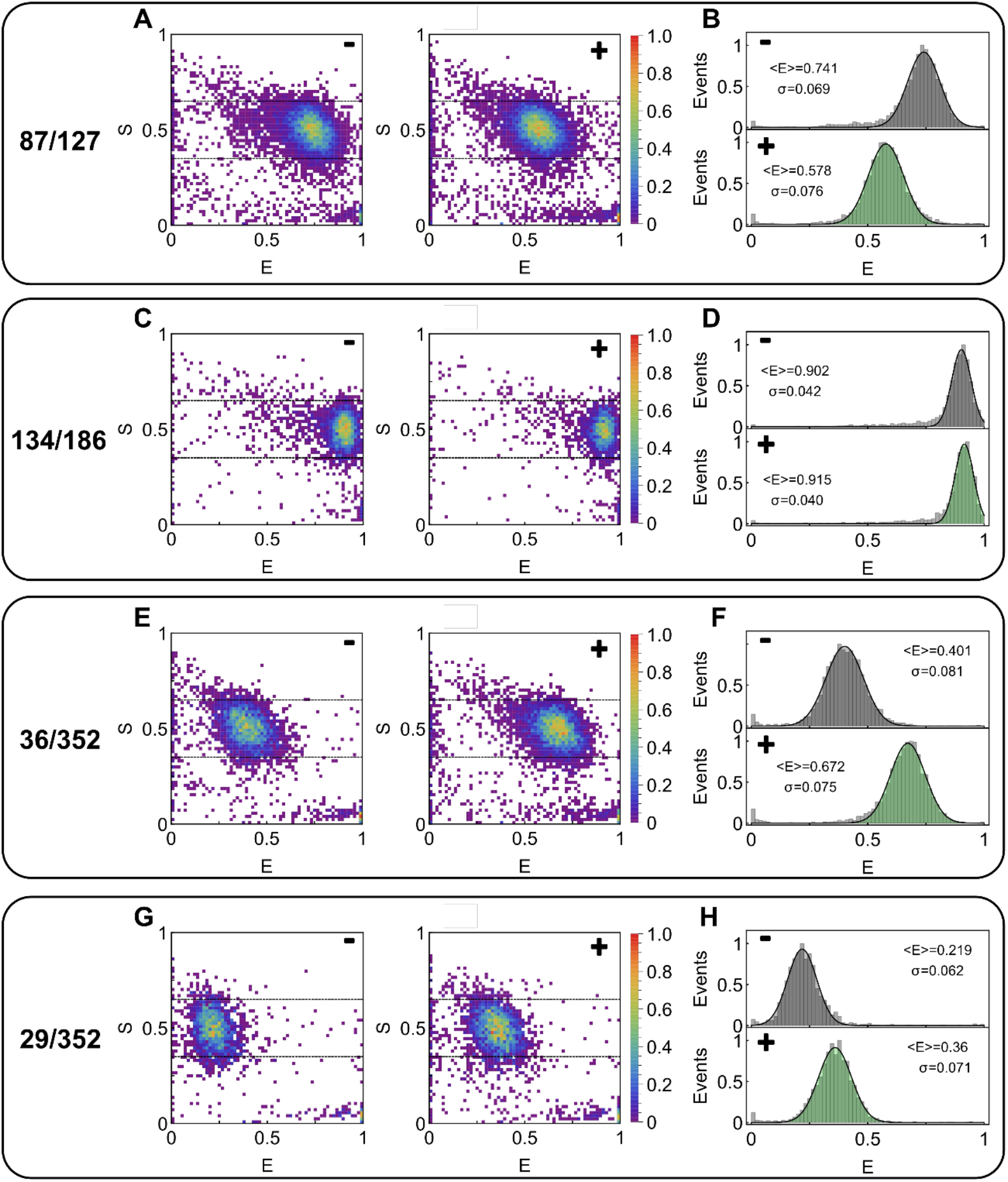
smFRET data of MalE with Alexa Fluor 555 – Alexa Fluor 647. **A, C, E, G)** ES-2D-Histograms of MalE mutants 87/186, 134/186, 36/352, and 29/352 in apo state (-) and holo state (+). **B, D, F, H)** 1D-E-Histograms extracted from the ES-Data for apo (grey) and holo (green) are fitted with a 1D-Gaussian distribution (). Mean <E> and standard deviation σ are labelled.

**SI Figure 9:**
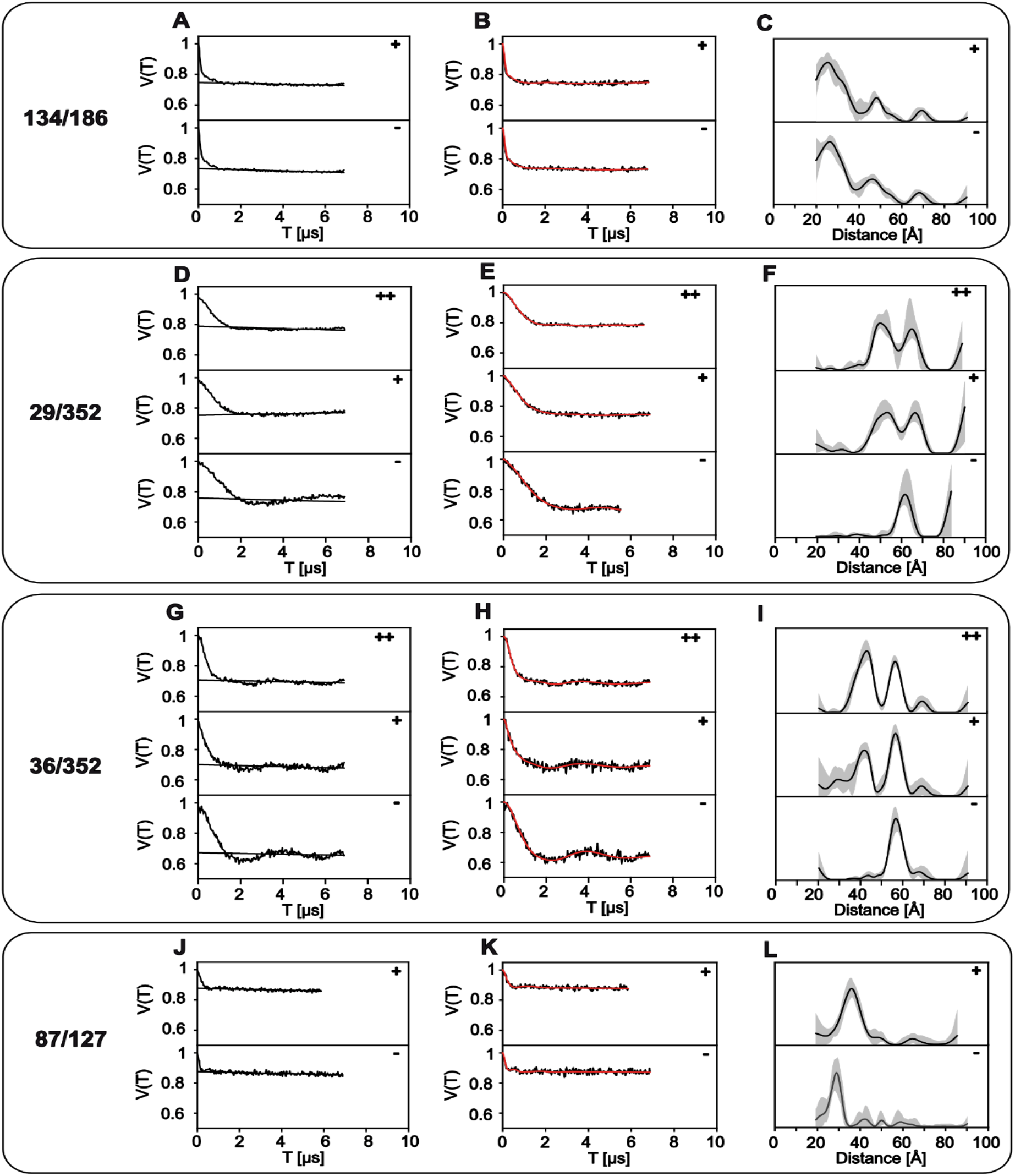
PELDOR/DEER data of MalE mutants from DEERanalysis. **A, D, G, J)** Raw PELDOR/DEER time traces for apo (-) and holo (+, 1 mM; ++, 10 mM maltose) measurements of each double mutant. The background, which was used for correction of the signal, is indicated as black line. **B, E, H, K)** Background-corrected PELDOR/DEER time traces (black) and fits of the signal (red). **C, F, I, L)** Distance distributions from PELDOR/DEER time traces (black) with validation of the distribution (grey).

**SI Figure 10:**
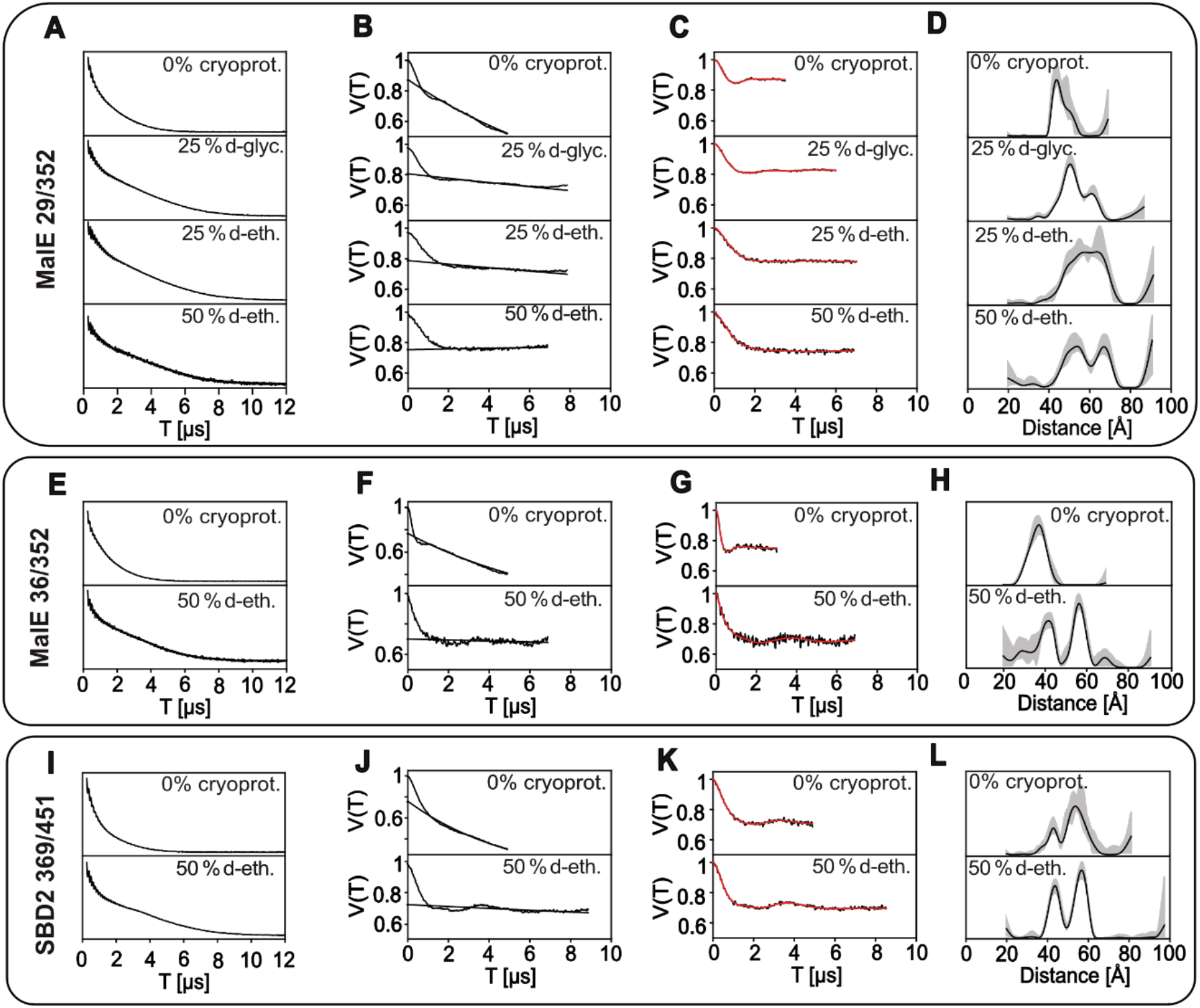
Data of PELDOR/DEER experiments with variations of cryoprotectant. **A, E, I)** 2PESEEM (2-pulse electron spin echo envelope modulation) spectra for each mutant with variations of cryoprotectant and the original 50% d-ethylene glycol measurements. **B, F, J)** Raw PELDOR/DEER time traces for each measurement. The background, which was used for correction of the signal, is indicated as black line. **C, G, K)** Background-corrected PELDOR/DEER time traces (black) and fits of the signal (red). **D, H, L)** Distance distributions from PELDOR/DEER time traces (black) with validation of the distribution (grey).

**SI Figure 11:**
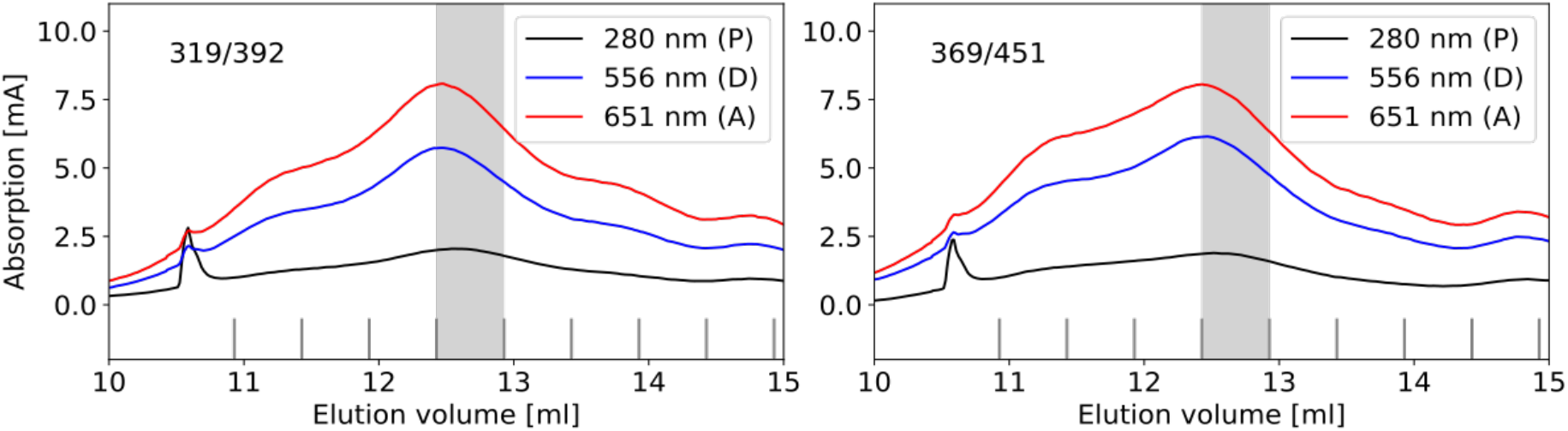
Size exclusion chromatography of SBD2 mutants labelled with Alexa Fluor 555 – Alexa Fluor 647. Absorption profile of the size extrusion chromatography (ÄKTA, Superdex 75 Increase 10/300 GL, GE Healthcare) for all tested SBD2 mutants 319/392 and 369/451 to monitor protein concentration (280 nm) and Alexa Fluor 555 (552 nm) / Alexa Fluor 647 (650 nm). The grey area indicates the fraction used in the smFRET experiments, where labelling efficiencies of >90% was achieved for all samples.

**SI Figure 12:**
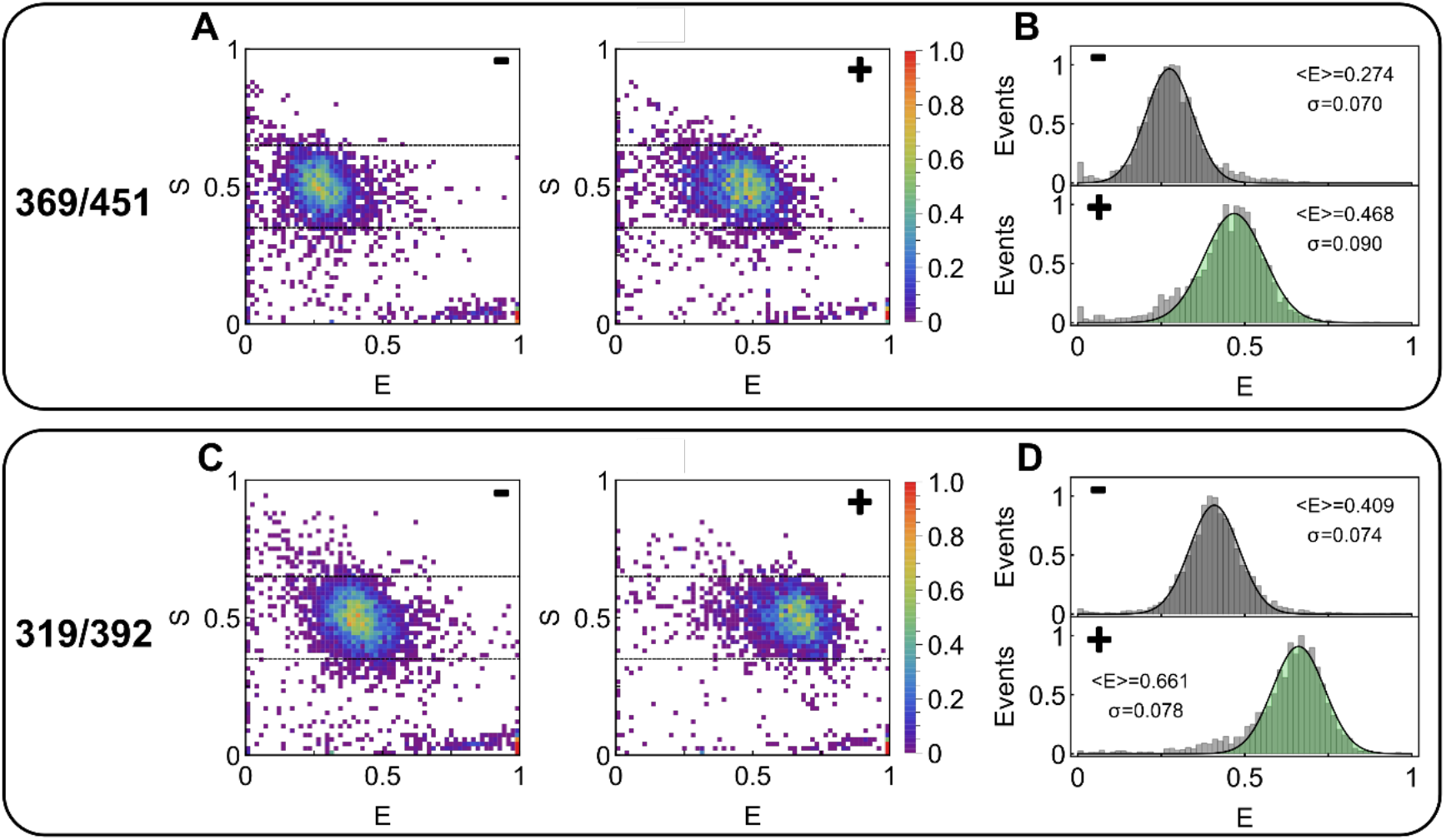
smFRET data of SBD2 with Alexa Fluor 555 – Alexa Fluor 647. **A, C)** ES-2D-Histograms of SBD2 mutants 369/451 and 319/392 in apo state (-) and holo state (+). **B, D)** 1D-E-Histograms extracted from the ES-Data for apo (grey) and holo (green) are fitted with a 1D-Gaussian distribution. Mean <E> and standard deviation σ are labelled.

**SI Figure 13:**
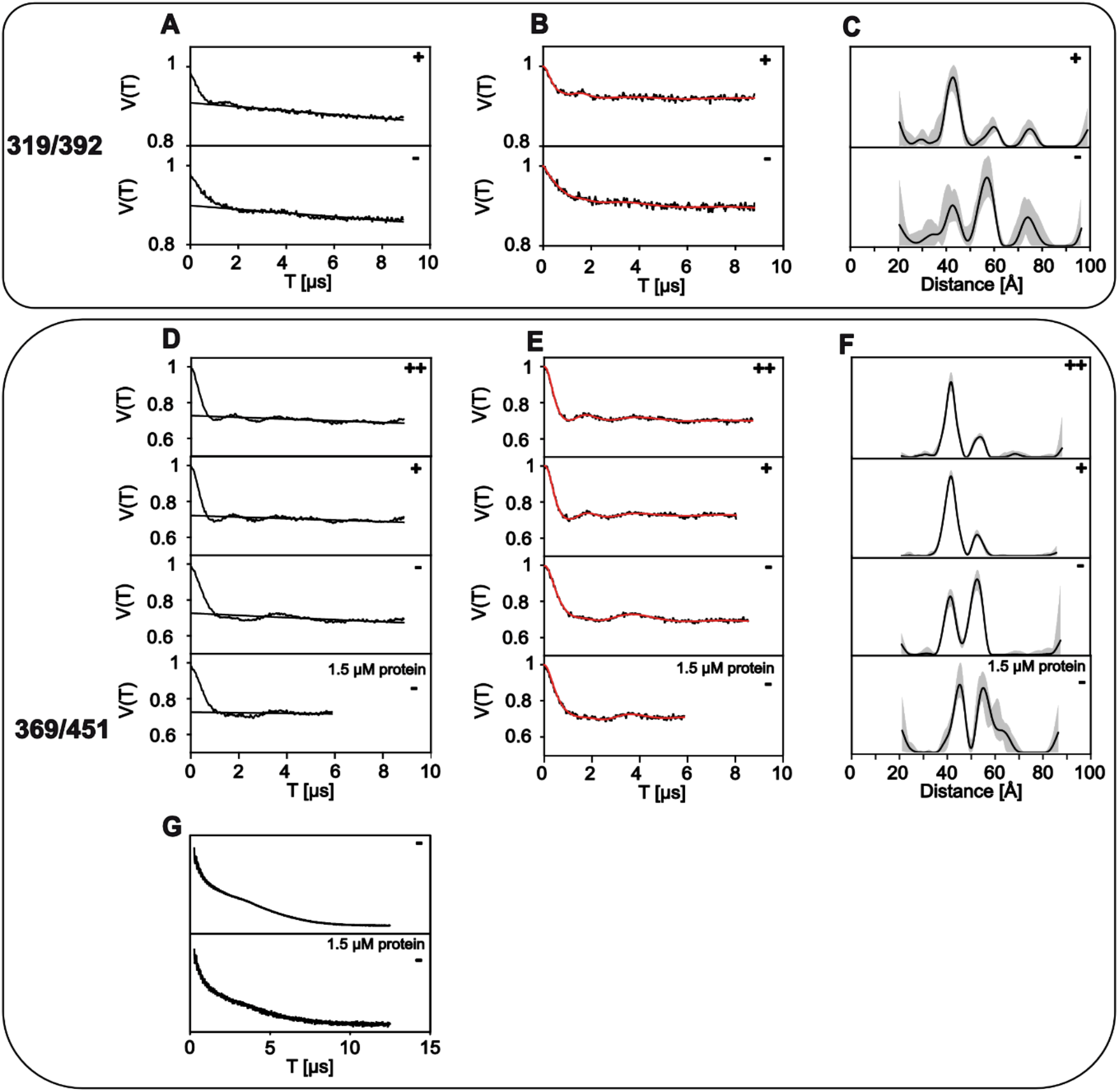
PELDOR/DEER data of SBD2 mutants from DEERanalysis. **A, D)** Raw PELDOR/DEER time traces for apo (-) and holo (+ and ++) measurements of each double mutant. The background, which was used for correction of the signal, is indicated as black line. **B, E)** Background-corrected PELDOR/DEER time traces (black) and fits of the signal (red). **C, F)** Distance distributions from PELDOR/DEER time traces (black) with validation of the distribution (grey). **G)** 2PESEEM spectra of apo measurements for 369/451 for 15 μM and 1.5 μM protein concentrations.

**SI Figure 14:**
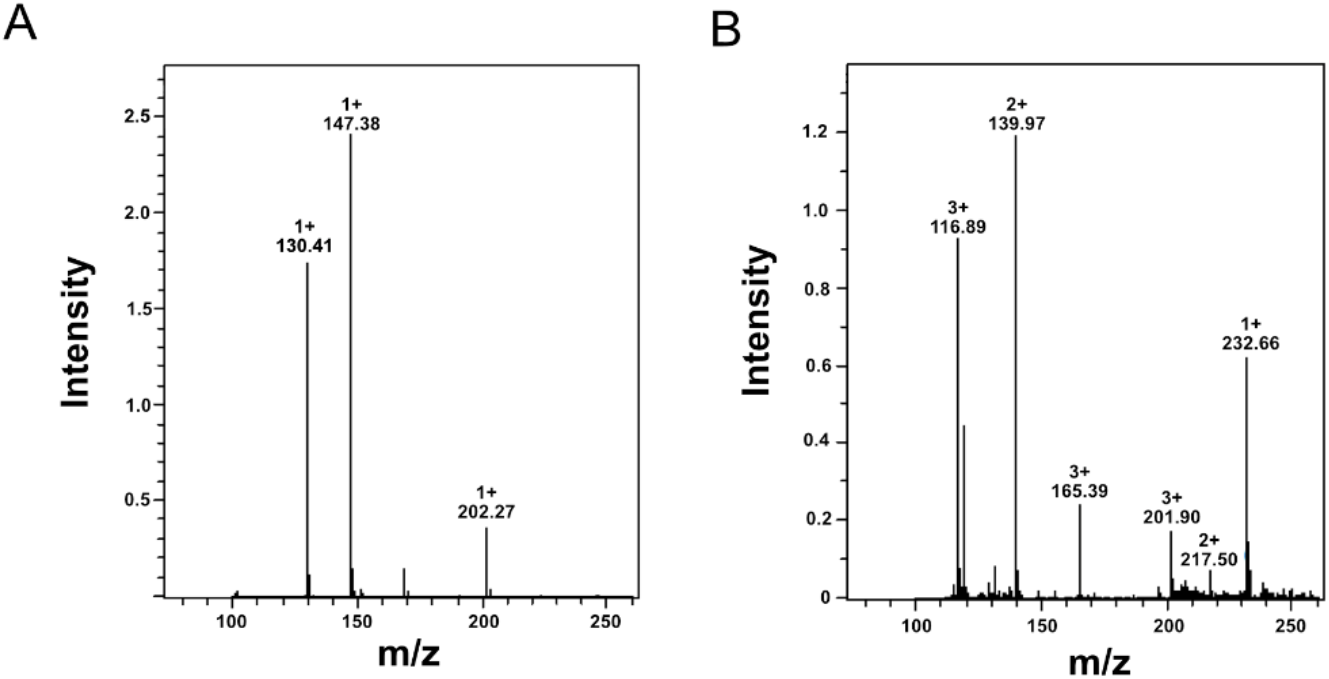
LC-MS for detection of glutamine. **A)** Mass spectrum of glutamine in standard protein buffer as positive control (MW_glutamine_: 146.15 g/mol). **B)** Mass spectrum of supernatant after precipitation and centrifugation of SBD2 protein sample.

**SI Figure 15:**
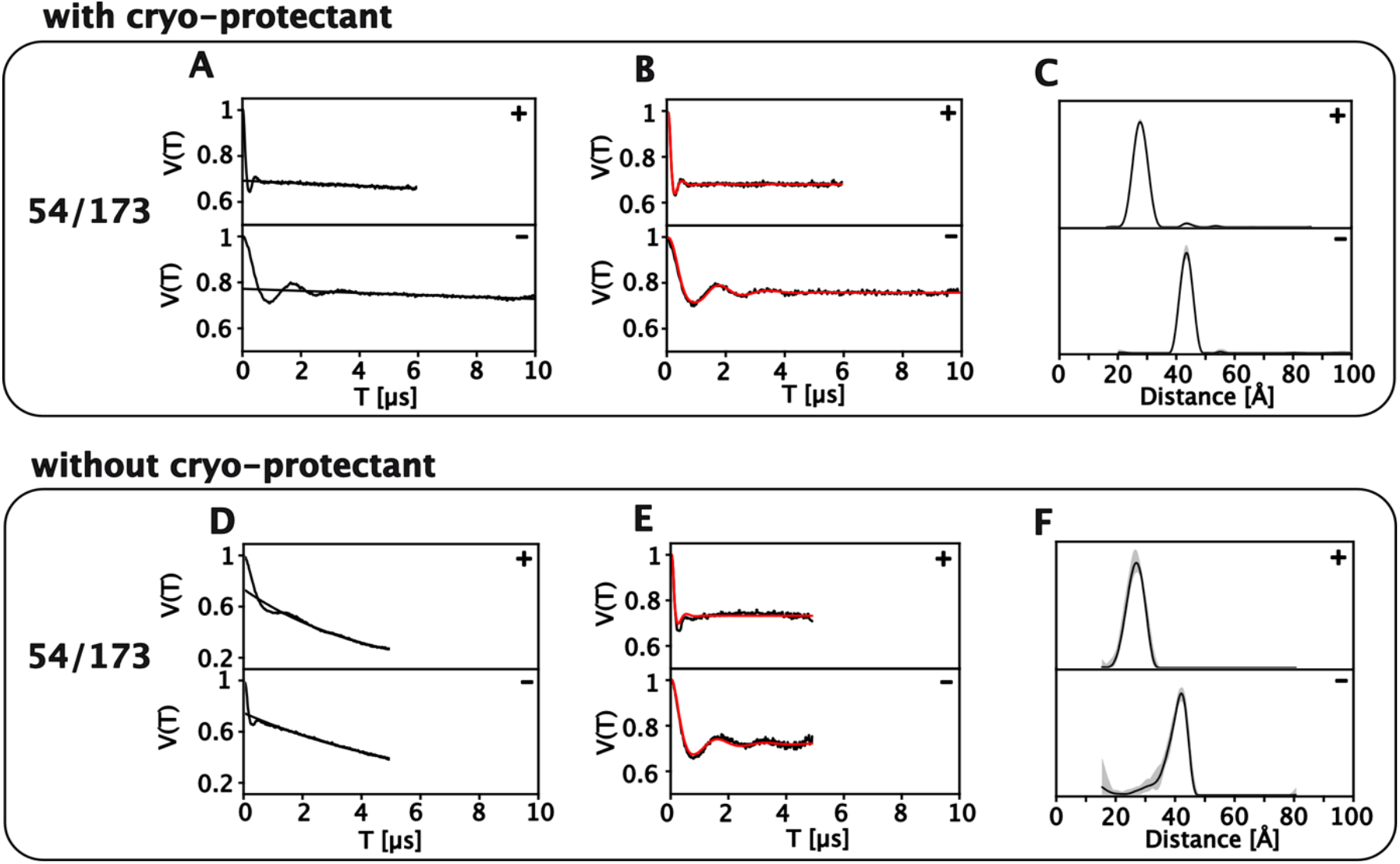
Comparison of PELDOR/DEER measurements on SiaP from *V. cholerae* with and without cryo-protectant. **A, D)** Raw PELDOR/DEER time traces for apo (-) and holo (+, 1 mM Neu5Ac) measurements of each double mutant. The background, which was used for correction of the signal, is indicated as black line. **B, E)** Background-corrected PELDOR/DEER time traces (black) and fits of the signal (red). **C, F)** Distance distributions from PELDOR/DEER time traces (black) with validation of the distribution (grey).

**SI Figure 16:**
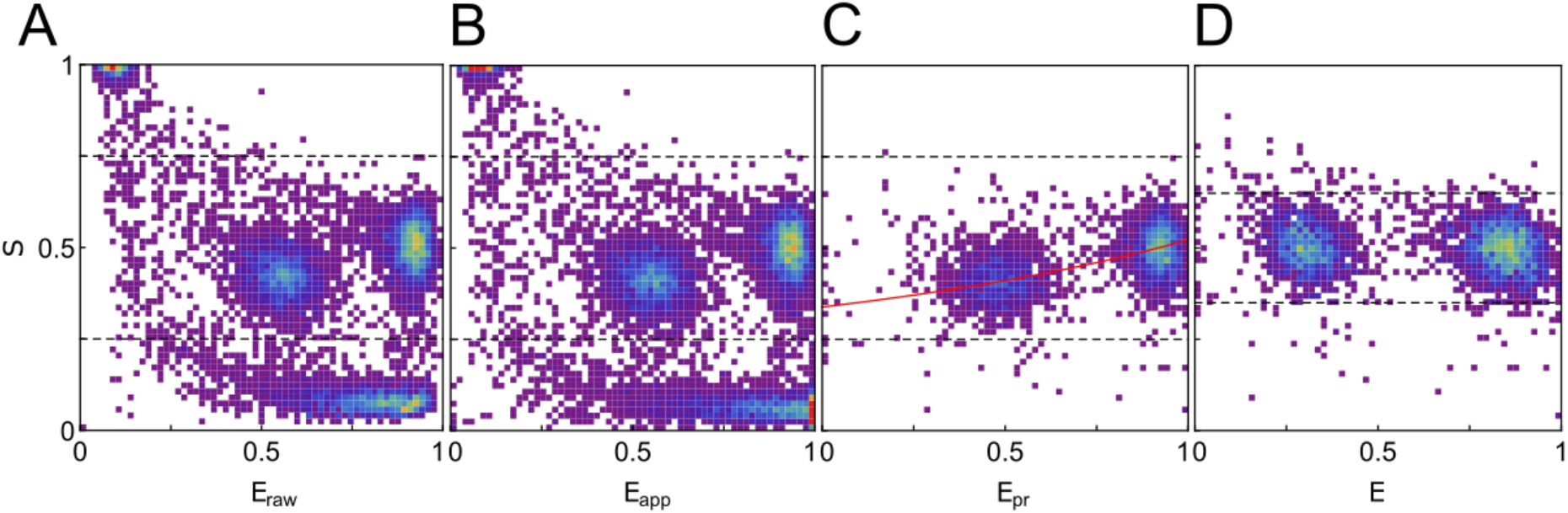
Correction procedure for smFRET to obtain setup-independent FRET efficiencies. **A)** Raw data in ES-histogram. **B)** Background corrected data. **C)**Removal of donor and acceptor only species and correction for spectral overlap of donor and acceptor fluorophore. The red line indicates the fit of the detection efficiency and quantum efficiency ratio of donor and detector, which leads to slope in S (along E). **D)** Correction for detection and quantum efficiencies.

## References

1. Locher, K. P. Structure and mechanism of ATP-binding cassette transporters. Philosophical Transactions of the Royal Society B: Biological Sciences 364, 239–245 (2009).

2. Thomas, C. & Tampé, R. Structural and mechanistic principles of ABC transporters. Annual review of biochemistry 89, 605–636 (2020).

3. Huse, M. & Kuriyan, J. The conformational plasticity of protein kinases. Cell 109, 275–282 (2002).

4. Loveridge, E.J., Behiry, E. M., Guo, J. & Allemann, R. K. Evidence that a ‘dynamic knockout’in Escherichia coli dihydrofolate reductase does not affect the chemical step of catalysis. Nature chemistry 4, 292–297 (2012).

5. Hofmann, S. e t al. Conformation space of a heterodimeric ABC exporter under turnover conditions. Nature 571, 580–583 (2019).

6. Wales, T. E. & Engen, J. R. Hydrogen exchange mass spectrometry for the analysis of protein dynamics. Mass spectrometry reviews 25, 158–170 (2006).

7. Förster, T. Zwischenmolekulare Energiewanderung und Fluoreszenz. Annalen der Physik 437, 55–75 (1948).

8. Ha, T. e t al. Probing the interaction between two single molecules: fluorescence resonance energy transfer between a single donor and a single acceptor. Proceedings of the National Academy of Sciences 93, 6264–6268 (1996).

9. Kapanidis, A. N. e t al. Fluorescence-aided molecule sorting: analysis of structure and interactions by alternating-laser excitation of single molecules. Proceedings of the National Academy of Sciences 101, 8936–8941 (2004).

10. Hohlbein, J., Craggs, T. D. & Cordes, T. Alternating-laser excitation: single-molecule FRET and beyond. Chemical Society Reviews 43, 1156–1171 (2014).

11. Tuukkanen, A. T., Spilotros, A. & Svergun, D. I. Progress in small-angle scattering from biological solutions at high-brilliance synchrotrons. IUCrJ 4, 518–528 (2017).

12. Jeschke, G. The contribution of modern EPR to structural biology. Emerging Topics in Life Sciences 2, ETLS20170143–18 (2018).

13. Müller, A. e t al. Conservation of structure and mechanism in primary and secondary transporters exemplified by SiaP, a sialic acid binding virulence factor from Haemophilus influenzae. Journal of Biological Chemistry 281, 22212–22222 (2006).

14. Mulligan, C., Fischer, M. & Thomas, G. H. Tripartite ATP-independent periplasmic (TRAP) transporters in bacteria and archaea. FEMS Microbiology Reviews 35, 68–86 (2011).

15. Hall, J. A., Gehring, K. & Nikaido, H. Two Modes of Ligand Binding in Maltose-binding Protein of Escherichia coli

16. Tang, C., Schwieters, C. D. & Clore, G. M. Open-to-closed transition in apo maltose-binding protein observed by paramagnetic NMR. Nature 449, 1078–1082 (2007).

17. Gouridis, G. e t al. Conformational dynamics in substrate-binding domains influences transport in the ABC importer GlnPQ. Nature Structural & Molecular Biology 22, 57 (2015).

18. Fulyani, F. e t al. Functional diversity of tandem substrate-binding domains in ABC transporters from pathogenic bacteria. Structure 21, 1879–1888 (2013).

19. Schuurman-Wolters, G. K. & Poolman, B. Substrate Specificity and Ionic Regulation of GlnPQ from Lactococcus lactis an ATP-binding cassette transporter with four extracytoplasmatic substrate-binding domains. Journal of Biological Chemistry 280, 23785–23790 (2005).

20. Van Der Velde, J. H. M. e t al. A simple and versatile design concept for fluorophore derivatives with intramolecular photostabilization. Nature communications 7, 10144 (2016).

21. Berntsson, R. P. A., Smits, S. H. J., Schmitt, L., Slotboom, D.-J. & Poolman, B. A structural classification of substrate-binding proteins. FEBS Letters 584, 2606–2617 (2010).

22. Phillips, G. N., Mahajan, V. K., Siu, A. K. & Quiocho, F. A. Structure of L-arabinose-binding protein from Escherichia coli at 5 A resolution and preliminary results at 3.5 A. Proceedings of the National Academy of Sciences 73, 2186–2190 (1976).

23. Quiocho, F. A. & Vyas, N. K. Novel stereospecificity of the L-arabinose-binding protein. Nature 310, 381–386 (1984).

24. Glaenzer, J., Peter, M. F., Thomas, G. H. & Hagelueken, G. PELDOR Spectroscopy Reveals Two Defined States of a Sialic Acid TRAP Transporter SBP in Solution. Biophysical Journal 112, 109–120 (2017).

25. Mao, B., Pear, M. R., McCammon, J. A. & Quiocho, F. A. Hinge-bending in L-arabinose-binding protein. The” Venus’s-flytrap” model. Journal of Biological Chemistry 257, 1131–1133 (1982).

26. Stockner, T., Vogel, H. J. & Tieleman, D. P. A salt-bridge motif involved in ligand binding and large-scale domain motions of the maltose-binding protein. Biophysical journal 89, 3362–3371 (2005).

27. Feng, Y. et al. Conformational Dynamics of apo-GlnBP Revealed by Experimental and Computational Analysis. Angewandte Chemie International Edition 55, 13990–13994 (2016).

28. de Boer, M., Gouridis, G., Muthahari, Y. A. & Cordes, T. Single-molecule observation of ligand binding and conformational changes in FeuA. Biophysical Journal 117, 1642–1654 (2019).

29. Kim, E. et al. A single-molecule dissection of ligand binding to a protein with intrinsic dynamics. Nature Chemical Biology 9, 313 (2013).

30. Husada, F. et al. Watching conformational dynamics of ABC transporters with single-molecule tools. Biochemical Society Transactions 43, 1041–1047 (2015).

31. de Boer, M. et al. Conformational and dynamic plasticity in substrate-binding proteins underlies selective transport in ABC importers. Elife 8, e44652 (2019).

32. Marinelli, F. et al. Evidence for an allosteric mechanism of substrate release from membrane-transporter accessory binding proteins. Proceedings of the National Academy of Sciences 108, E1285–92 (2011).

33. Marinelli, F. & Fiorin, G. Structural Characterization of Biomolecules through Atomistic Simulations Guided by DEER Measurements. Structure 27, 359–370.e12 (2019).

34. Quiocho, F. A., Spurlino, J. C. & Rodseth, L. E. Extensive features of tight oligosaccharide binding revealed in high-resolution structures of the maltodextrin transport/chemosensory receptor. Structure 5, 997–1015 (1997).

35. Johnston, J. W. et al. Characterization of the N-acetyl-5-neuraminic acid-binding site of the extracytoplasmic solute receptor (SiaP) of nontypeable Haemophilus influenzae strain 2019. Journal of Biological Chemistry 283, 855–865 (2008).

36. Gangi Setty, T., Cho, C., Govindappa, S., Apicella, M. A. & Ramaswamy, S. Bacterial periplasmic sialic acid-binding proteins exhibit a conserved binding site. Acta Crystallographica Section D 70, 1801–1811 (2014).

37. Burger, M., Rein, S., Weber, S., Gräber, P. & Kacprzak, S. Distance measurements in the F0F1-ATP synthase from E. coli using smFRET and PELDOR spectroscopy. European Biophysics Journal 1–10 (2019).

38. Klose, D. et al. Simulation vs. reality: a comparison of in silico distance predictions with DEER and FRET measurements. PLoS one 7, e39492 (2012).

39. Grohmann, D. et al. RNA-binding to archaeal RNA polymerase subunits F/E: a DEER and FRET study. Journal of the American Chemical Society 132, 5954–5955 (2010).

40. Jeschke, G., Chechik, V., Ionita, P. & Godt, A. DeerAnalysis2006—a comprehensive software package for analyzing pulsed ELDOR data. Applied Magnetic Resonance 30, 473–498 (2006).

41. Glaenzer, J., Peter, M. F. & Hagelueken, G. Studying structure and function of membrane proteins with PELDOR/DEER spectroscopy - The crystallographers perspective. Methods (San Diego, Calif) 147, 163–175 (2018).

42. Jeschke, G. & Polyhach, Y. Distance measurements on spin-labelled biomacromolecules by pulsed electron paramagnetic resonance. Physical Chemistry Chemical Physics 9, 1895–1910 (2007).

43. Borbat, P. P. & Freed, J. H. Pulse dipolar electron spin resonance: distance measurements. Structural Information from Spin-Labels and Intrinsic Paramagnetic Centres in the Biosciences (2013).

44. Kulik, L. V., Dzuba, S. A., Grigoryev, I. A. & Tsvetkov, Y. D. Electron dipole–dipole interaction in ESEEM of nitroxide biradicals. 343, 315–324 (2001).

45. Jeschke, G., Pannier, M., Godt, A. & Spiess, H. W. Dipolar spectroscopy and spin alignment in electron paramagnetic resonance. Chemical Physics Letters 331, 243–252 (2000).

46. Jeschke, G. DEER Distance Measurements on Proteins. Annual Review of Physical Chemistry 63, 419–446 (2012).

47. Schiemann, O. & Prisner, T. F. Long-range distance determinations in biomacromolecules by EPR spectroscopy. Quarterly Reviews of Biophysics 40, 1–53 (2007).

48. Klare, J. P. & Steinhoff, H. J. Spin labeling EPR. Photosynthesis Research 102, 377–390 (2009).

49. Dimura, M. et al. Quantitative FRET studies and integrative modeling unravel the structure and dynamics of biomolecular systems. Current Opinion in Structural Biology 40, 163–185 (2016).

50. Galazzo, L. et al. Spin-labeled nanobodies as protein conformational reporters for electron paramagnetic resonance in cellular membranes. Proceedings of the National Academy of Sciences 117, 2441–2448 (2020).

51. Fleissner, M. R. et al. Site-directed spin labeling of a genetically encoded unnatural amino acid. Proceedings of the National Academy of Sciences of the United States of America 106, 21637–21642 (2009).

52. Schmidt, M. J., Borbas, J., Drescher, M. & Summerer, D. A genetically encoded spin label for electron paramagnetic resonance distance measurements. Journal of the American Chemical Society 136, 1238–1241 (2014).

53. Roy, R., Hohng, S. & Ha, T. A practical guide to single-molecule FRET. Nature methods 5, 507–516 (2008).

54. Lee, T. C. et al. Dual unnatural amino acid incorporation and click-chemistry labeling to enable single-molecule FRET studies of p97 folding. Chembiochem: a European journal of chemical biology 17, 981 (2016).

55. El Mkami, H. & Norman, D. G. 125–152 (Elsevier, 2015).

56. Schmidt, T., Wälti, M. A., Baber, J. L., Hustedt, E. J. & Clore, G. M. Long Distance Measurements up to 160 Å in the GroEL Tetradecamer Using Q-Band DEER EPR Spectroscopy. Angewandte Chemie (International ed in English) (2016).

57. Krainer, G., Hartmann, A. & Schlierf, M. farFRET: extending the range in single-molecule FRET experiments beyond 10 nm. Nano letters 15, 5826–5829 (2015).

58. Jeschke, G. MMM: A toolbox for integrative structure modeling. Protein Science 181, 223–285 (2017).

59. Hagelueken, G., Abdullin, D., Ward, R. & Schiemann, O. mtsslSuite: In silico spin labelling, trilateration and distance-constrained rigid body docking in PyMOL. Molecular Physics 111, 2757–2766 (2013).

60. Kalinin, S. et al. A toolkit and benchmark study for FRET-restrained high-precision structural modeling. Nature Methods 9, 1218–1225 (2012).

61. Muschielok, A. et al. A nano-positioning system for macromolecular structural analysis. Nature Methods 5, 965–971 (2008).

62. Beckers, M., Drechsler, F., Eilert, T., Nagy, J. & Michaelis, J. Quantitative structural information from single-molecule FRET. Faraday discussions 184, 117–129 (2015).

63. Gebhardt, C., Lehmann, M., Reif, M., Zacharias, M. & Cordes, T. Molecular and spectroscopic characterization of green and red cyanine fluorophores from the Alexa Fluor and AF series. bioRxiv (2020).

64. Sauer, M., Hofkens, J. & Enderlein, J. Handbook of fluorescence spectroscopy and imaging: from ensemble to single molecules (John Wiley & Sons, 2010).

65. Berliner, L. J., Grunwald, J., Hankovszky, H. O. & Hideg, K. A novel reversible thiol-specific spin label: papain active site labeling and inhibition. Analytical Biochemistry 119, 450–455 (1982).

66. Yagi, H. et al. Gadolinium tagging for high-precision measurements of 6 nm distances in protein assemblies by EPR. 133, 10418–10421 (2011).

67. Reginsson, G. W., Kunjir, N. C., Sigurdsson, S. T. & Schiemann, O. Trityl Radicals: Spin Labels for Nanometer-Distance Measurements. Chemistry - A European Journal 18, 13580–13584 (2012).

68. Hagelueken, G., Ward, R., Naismith, J. H. & Schiemann, O. MtsslWizard: In Silico Spin-Labeling and Generation of Distance Distributions in PyMOL. Applied Magnetic Resonance 42, 377–391 (2012).

69. Hagelueken, G., Abdullin, D. & Schiemann, O. mtsslSuite: Probing Biomolecular Conformation by Spin-Labeling Studies. Methods in Enzymology 563, 595–622 (2015).

70. Kalinin, S. et al. A toolkit and benchmark study for FRET-restrained high-precision structural modeling. Nature methods 9, 1218–1225 (2012).

71. Hellenkamp, B. et al. Precision and accuracy of single-molecule FRET measurements-a multi-laboratory benchmark study. Nature Methods 15, 669–676 (2018).

72. Kudryavtsev, V. et al. Combining MFD and PIE for accurate single-pair Förster resonance energy transfer measurements. ChemPhysChem 13, 1060–1078 (2012).

73. Peulen, T.-O., Opanasyuk, O. & Seidel, C. A. M. Combining graphical and analytical methods with molecular simulations to analyze time-resolved FRET measurements of labeled macromolecules accurately. The Journal of Physical Chemistry B 121, 8211–8241 (2017).

74. Ploetz, E. et al. Förster resonance energy transfer and protein-induced fluorescence enhancement as synergetic multi-scale molecular rulers. Scientific reports (2016).

75. Lillington, J. E. D. et al. Shigella flexneri Spa15 crystal structure verified in solution by double electron electron resonance. Journal of Molecular Biology 405, 427–435 (2011).

76. Hatmal, M. M. et al. Computer modeling of nitroxide spin labels on proteins. Biopolymers 97, 35–44 (2011).

77. Bordenave, T. et al. Synthesis and in vitro and in vivo evaluation of MMP-12 selective optical probes. Bioconjugate chemistry 27, 2407–2417 (2016).

78. Fleissner, M. R. et al. Structure and dynamics of a conformationally constrained nitroxide side chain and applications in EPR spectroscopy. Proceedings of the National Academy of Sciences of the United States of America 108, 16241–16246 (2011).

79. Fleck, N. et al. SLIM: A Short-Linked, Highly Redox-Stable Trityl Label for High-Sensitivity In-Cell EPR Distance Measurements. Angewandte Chemie (2020).

80. Potapov, A. et al. Nanometer-scale distance measurements in proteins using Gd3+ spin labeling. 132, 9040–9048 (2010).

81. Cunningham, T. F., Putterman, M. R., Desai, A., Horne, W. S. & Saxena, S. The Double-Histidine Cu 2+-Binding Motif: A Highly Rigid, Site-Specific Spin Probe for Electron Spin Resonance Distance Measurements. Angewandte Chemie 127, 6428–6432 (2015).

82. Alexander, N. S. et al. RosettaEPR: rotamer library for spin label structure and dynamics. PLoS one 8, e72851 (2013).

83. Jeschke, G. Conformational dynamics and distribution of nitroxide spin labels. Progress in nuclear magnetic resonance spectroscopy 72, 42–60 (2013).

84. Spicher, S. & Grimme, S. Robust Atomistic Modeling of Materials, Organometallic, and Biochemical Systems. Angewandte Chemie (2020).

85. Spicher, S., Abdullin, D., Grimme, S. & Schiemann, O. Modeling of spin-spin distance distributions for nitroxide labeled biomacromolecules. Physical Chemistry Chemical Physics (2020).

86. Schmidt, T., Jeon, J., Okuno, Y., Chiliveri, S. C. & Clore, G. M. Sub-millisecond freezing coupled permits cryoprotectant-free EPR double electron-electron resonance spectroscopy. ChemPhysChem (2020).

87. Vagenende, V., Yap, M. G. S. & Trout, B. L. Mechanisms of protein stabilization and prevention of protein aggregation by glycerol. Biochemistry 48, 11084–11096 (2009).

88. Georgieva, E. R. et al. Effect of freezing conditions on distances and their distributions derived from Double Electron Electron Resonance (DEER): a study of doubly-spin-labeled T4 lysozyme. Journal of Magnetic Resonance 216, 69–77 (2012).

89. Vera, L. & Stura, E. A. Strategies for Protein Cryocrystallography. Crystal Growth & Design 14, 427–435 (2014).

90. Peter, M. F. et al. Studying Conformational Changes of the Yersinia Type-III-Secretion Effector YopO in Solution by Integrative Structural Biology. Structure (2019).

91. Alonso-García, N. et al. Combination of X-ray crystallography, SAXS and DEER to obtain the structure of the FnIII-3,4 domains of integrin α6β4. Acta Crystallographica Section D 71, 969–985 (2015).

92. Igarashi, R. et al. Distance determination in proteins inside Xenopus laevis oocytes by double electron-electron resonance experiments. Journal of the American Chemical Society 132, 8228–8229 (2010).

93. Jahromy, Y. N. & Schubert, E. Demystifying EPR: A Rookie Guide to the Application of Electron Paramagnetic Resonance Spectroscopy on Biomolecules. Progress in Biological Sciences (2014).

94. Kapanidis, A. N. et al. Fluorescence-aided molecule sorting: analysis of structure and interactions by alternating-laser excitation of single molecules. Proceedings of the National Academy of Sciences 101, 8936–8941 (2004).

95. Eggeling, C., Fries, J. R., Brand, L., Günther, R. & Seidel, C. A. M. Monitoring conformational dynamics of a single molecule by selective fluorescence spectroscopy. Proceedings of the National Academy of Sciences 95, 1556–1561 (1998).

96. Lee, N. K. et al. Accurate FRET measurements within single diffusing biomolecules using alternating-laser excitation. Biophysical journal 88, 2939–2953 (2005).

97. Torella, J. P., Holden, S. J., Santoso, Y., Hohlbein, J. & Kapanidis, A. N. Identifying molecular dynamics in single-molecule FRET experiments with burst variance analysis. Biophysical journal 100, 1568–1577 (2011).

98. Lakowicz, J. R. Principles of fluorescence spectroscopy (Springer science & business media, 2013).

99. Tsukanov, R., Tomov, T. E., Berger, Y., Liber, M. & Nir, E. Conformational dynamics of DNA hairpins at millisecond resolution obtained from analysis of single-molecule FRET histograms. The Journal of Physical Chemistry B 117, 16105–16109 (2013).

